# *Cis*-vaccenic acid is a key product of stearoyl-CoA desaturase 1 and a critical oncogenic factor in prostate cancer

**DOI:** 10.1101/2024.03.28.587238

**Authors:** Julia S Scott, Reuben SE Young, Lake-Ee Quek, Deanna C Miller, Emma Evergren, Jonas Dehairs, Ian RD Johnson, Doug A Brooks, Massimo Loda, Andrew J Hoy, Stephen J Blanksby, Johannes V Swinnen, Zeyad D Nassar, Lisa M Butler

**Affiliations:** South Australian immunoGENomics Cancer Institute, Adelaide Medical School and Freemasons Centre for Male Health and Wellbeing, University of Adelaide, Adelaide SA 5005, Australia; Precision Cancer Medicine Theme, South Australian Health and Medical Research Institute, Adelaide, SA 5000, Australia; Mass Spectrometry Development Laboratory, Queensland University of Technology, Brisbane, Qld, Australia; School of Chemistry and Molecular Bioscience, University of Wollongong, Wollongong, NSW, Australia; School of Mathematics and Statistics, The University of Sydney, Camperdown, NSW, Australia; Patrick G Johnston Centre for Cancer Research, Queen’s University Belfast, Belfast, UK; LKI – Leuven Cancer Institute, Department of Oncology, Laboratory of Lipid Metabolism and Cancer, KU Leuven, Leuven, B-3000, Belgium; Clinical and Health Sciences, University of South Australia, Adelaide, South Australia 5000, Australia; Department of Pathology and Laboratory Medicine, Weill Cornell Medicine, New York-Presbyterian Hospital, New York, New York; School of Medical Sciences, Charles Perkins Centre, Faculty of Medicine and Health, The University of Sydney, Sydney, New South Wales, Australia

**Keywords:** monounsaturated fatty acids, prostate cancer, cardiolipins, *cis*-vaccenic acid, fatty acid metabolism

## Abstract

Altered monounsaturated fatty acid (MUFA) metabolism is a hallmark of oncogenic transformation. Recently, the MUFA *cis*-vaccenic acid (cVA) was identified as a putative regulator of prostate cancer cell viability. cVA production requires the activity of stearoyl CoA desaturase 1 (SCD1), an enzyme frequently dysregulated in cancer, but the role of cVA in regulating cancer cell phenotypes has not been extensively explored, compared with the more commonly studied product and structural isomer of cVA, oleic acid. Here, we utilised SCD1 inhibition to study the effects of cVA in prostate cancer cells. We found that cVA consistently rescues reductions in cell viability due to SCD1 inhibition and promotes cell growth under normal conditions, thereby identifying cVA as an important and previously unrecognised product of SCD1 in prostate cancer. More broadly, we demonstrate that individual MUFA species exert a diverse range of influence on oncogenic phenotypes, highlighting a need to more precisely characterise the lipidome of cancer cells to understand the molecular pathology of the disease.

## Introduction

Altered lipid metabolism is a well-recognised feature of oncogenic transformation^1^, hijacked by cancer cells to promote cell growth and proliferation^2,3^. This is well-studied in prostate cancer, which displays increased reliance on fatty acid oxidation as the predominant energy source compared to other solid malignancies^4^. Furthermore, the expression of many key lipid metabolism genes is controlled by androgen signalling, the hormonal driver of disease, including fatty acid synthase (*FASN*) and stearoyl CoA desaturase 1 (*SCD1*)^5^, supporting targeting of lipid metabolism in prostate cancer as a promising therapeutic avenue.

We recently identified the fatty acid elongase enzyme, elongation of very long chain fatty acid 5 (ELOVL5) as a critical androgen-regulated oncogenic factor in prostate cancer cells^6^, required for prostate cancer cell viability due to the production of the monounsaturated fatty acid (MUFA), *cis*-vaccenic acid (cVA; FA 18:1*n*-7, *cis*)^6^. Mechanistically, ELOVL5 protein levels were positively correlated with mitochondrial respiration, and were inversely correlated with reactive oxygen species (ROS) production^6^, suggesting a link between ELOVL5, cVA and mitochondrial homeostasis. This finding is significant, as cVA is an often overlooked product of ELOVL5, compared to the polyunsaturated fatty acid (PUFA) species, arachidonic acid and docosahexaenoic acid^7^. cVA is an interesting fatty acid that is found in human milk and is linked with maternal obesity^8,9^, and regulation of lipotoxicity in *Saccharomyces cerevisiae* (baker’s yeast)^10^.

Increased MUFA abundance has been observed in a range of cancer types^11–13^, including prostate cancer^14^, and SCD1, the enzyme responsible for canonical MUFA production, is commonly upregulated^15,16^. In many *in vitro* studies, the effects of SCD1 inhibition are attributed to reduced production of oleic acid (OA; FA 18:1*n*-9, *cis*), an abundant product of SCD1^17^ (Fig. 1A), as supplementation of OA can rescue SCD1 inhibition-induced alterations to cell phenotypes^16,18,19^. SCD1 is also required for the production of cVA, as it converts palmitate (FA 16:0) to palmitoleic acid (FA 16:1*n*-7), which is subsequently elongated by ELOVL5 to generate cVA^20,21^ (Fig. 1A). Measurement of the relative abundance of cVA and OA – carried by membrane phospholipids – within prostate tissue suggest a strong association of OA with premalignant tissue, while cVA abundance correlated with immune cell infiltration and activation^22^. Indeed, palmitoleic acid has previously been shown to have lipokine activity^23^, with distinctive roles being demonstrated in cardiovascular disease, diabetes, obesity and hepatosteatosis^24^. Despite these previous findings, the elongation product of palmitoleic acid, cVA, has been understudied in the broader context of health and disease. Considering the high abundance of cVA in prostate cancer cell lines and tissues^25^, the specific role this fatty acid plays in oncogenesis and tumour progression is yet to be explored. Here, we utilised both SCD1 inhibition and ELOVL5 targeting to elucidate the role of cVA in prostate cancer more precisely, by studying prostate cancer cellular responses to broad MUFA depletion, specific cVA reduction, and individual MUFA supplementation, comparing the responses of cVA with OA.

**Figure 1.**
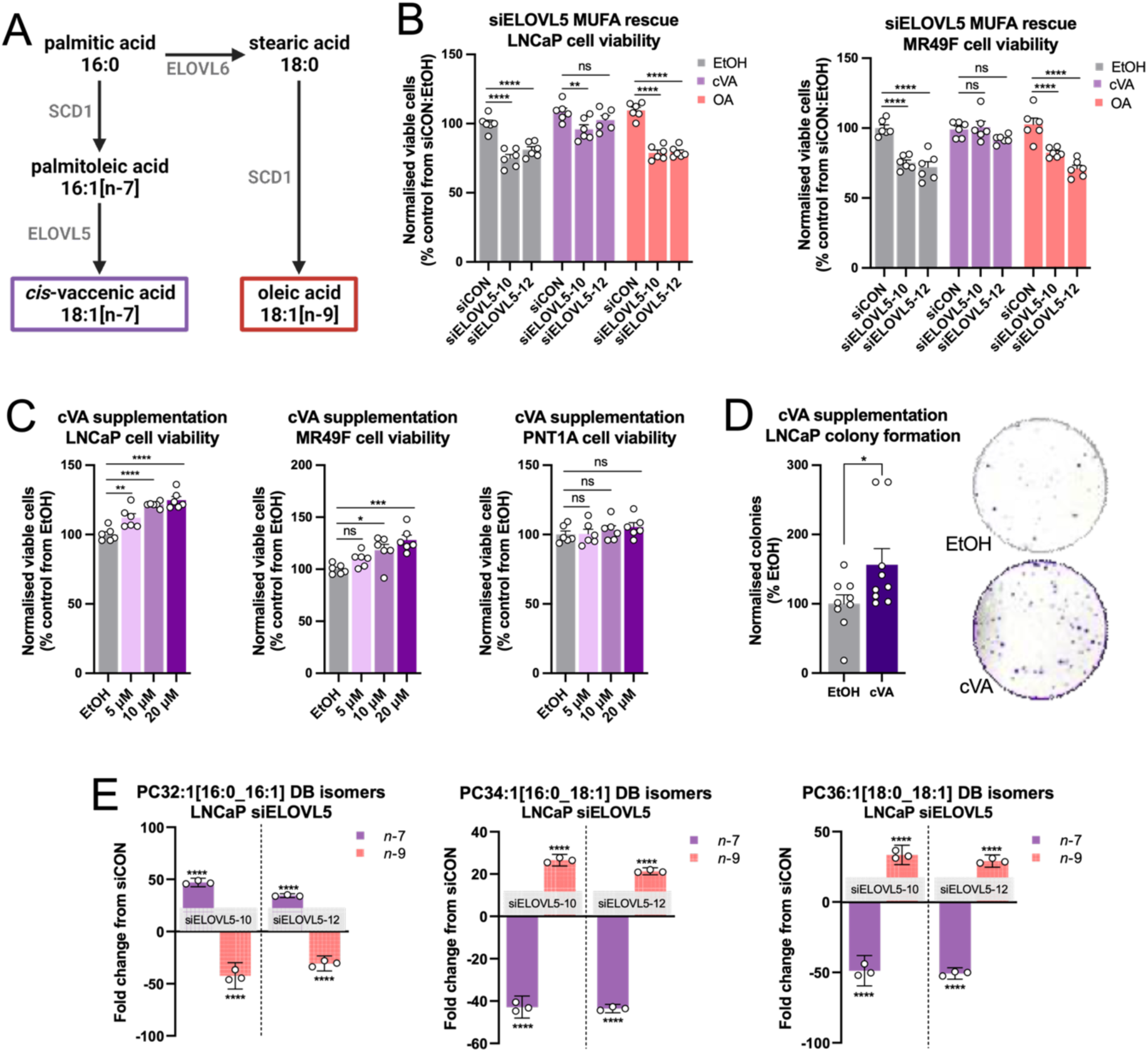
The MUFA product of ELOVL5, *cis*-vaccenic acid, promotes prostate cancer cell viability. [A] Schematic of *cis*-vaccenic acid (cVA; 18:1*n*-7, *cis*) and oleic acid (OA; 18:1*n*-9, *cis*) production. [B] Cell viability experiments under ELOVL5 knockdown (siELOVL5) and monounsaturated fatty acid (MUFA) supplementation of 10 µM cVA or 10 µM OA in prostate cancer cell lines LNCaP and MR49F, 96 hours post-transfection. Data are presented as normalised mean ± SEM of two independent experiments (N = 2), with three technical replicates (n = 3). [C] Cell viability measurements with cVA supplementation in LNCaP and MR49F prostate cancer cell lines, and benign prostate cell line PNT1A, 144 hours post-supplementation. Data are presented as normalised mean ± SEM (N = 2; n = 3). [D] Colony formation was evaluated with 100 µM cVA supplementation in LNCaP prostate cancer cell line, 2 weeks post-supplementation. Data are presented as mean ± SEM (N = 3; n = 3). [E] Ozone-induced dissociation (OzID) lipidomic analysis of *n*-7 and *n*-9 and isomer abundance in phosphatidylcholines (PC) with compositions 32:1, 34:1 and 36:1 following ELOVL5 knockdown (siELOVL5) in LNCaP cells, 96 hours post-transfection. Data are presented as a change in the relative abundance of three isomers with mean ± 95% confidence interval (CI) (N = 1; n = 3). Statistical significance was determined using two-way ANOVA and multiple comparisons [B & E], one-way ANOVA and multiple comparisons [C], or two-tailed student t-tests [D]. * p<0.05; ** p<0.01; *** p<0.001; ****p<0.0001.

## Materials and methods

### Cell culture

The human non-malignant prostate epithelial cell line PNT1A was obtained from the European Collection of Authenticated Cell Cultures (ECACC; England, UK), and the human prostate cancer cell lines LNCaP and 22Rv1 were purchased from the American Type Cell Collection (ATCC). Castrate-resistant and enzalutamide-resistant cell lines, V16D and MR49F, were derived from serial xenograft passages of LNCaP cell lines^26^ and were a kind gift from Professor Amina Zoubeidi’s laboratory (Vancouver Prostate Centre, BC, Canada). Human colorectal cancer cell line LOVO was generously provided by Associate Professor Susan Woods (The University of Adelaide, SA, Australia). All prostate cancer cell lines were authenticated by short tandem repeat profiling at Cell Bank Australia (NSW, Australia) in July 2020 and July 2022. Mycoplasma testing was carried out every 3-6 months to ensure cell line purity. Cells were cultured in RPMI-1640 medium (Sigma-Aldrich) containing 2 mmol/L L-glutamine (Sigma-Aldrich) and 10% foetal bovine serum (FBS) (Sigma-Aldrich; Cell Sera) and 10 µM enzalutamide (Cayman Chemical) for enzalutamide-resistant cell lines (MR49F and V16D). All chemicals were purchased from Sigma-Aldrich, unless otherwise stated. *Cis*-vaccenic acid (FA 18:1*n*-7, *cis*), palmitoleic acid (FA 16:1*n*-7, *cis*) and CAY10566 were purchased from Cayman Chemical. Isotopically labelled palmitoleic acid (U-13C16 98%) and oleic acid (U-13C18 98%) were purchased from Cambridge Isotope Laboratories. A939572 was purchased from Tocris Bioscience. Venetoclax was purchased from Abcam. *Cis*-vaccenic acid, palmitoleic acid, oleic acid (FA 18:1*n*-9, *cis*), ^13^C_16_-palmitoleic acid and ^13^C_18_-oleic acid were dissolved in ethanol. Enzalutamide, A939572, CAY10566, erastin, venetoclax and elesclomol were dissolved in DMSO. L-ascorbic acid was dissolved in nuclease-free water. The sources for the primary antibodies used in this study are listed in Supplementary Table S1.

### Transient RNA interference

#### Reverse transfection

ON-TARGET plus human small interfering RNAs (siRNAs) and a control siRNA (D-001810-01-20 ON-TARGET plus Nontargeting siRNA #1) were from Dharmacon (Millennium Science). The most effective siRNAs from a pool of four were selected for targeting ELOVL5 (J-009260-10, J-009260-12) and SCD1 (LQ-005061-00-0002), and reversed transfected at a concentration of 5 nmol/L using Lipofectamine RNAiMAX transfection reagent (Invitrogen) according to the manufacturer’s instructions.

#### Nucleofection

ON-TARGET plus human small interfering RNAs targeting ELOVL5 (J-009260-10, J-009260-12) and a control siRNA (D-001810-01-20 ON-TARGET plus Nontargeting siRNA #1) were prepared for electroporation using the Lonza SF Cell Line 4D-Nucleofector™ X Kit (V4XC-2012; Lonza) according to the manufacturer’s instruction with minor alterations. LNCaP cells were cultured until 80% confluent, before being harvested by trypsinisation and counted to determine cell density. The required number of cells (5x10^6^ per replicate) was centrifuged at 90 x *g* for 10 mins, and then resuspended in 4D-Nucleofector™ solution (100 µL per replicate), prepared according to the manufacturer’s instructions. The cell suspension was divided into the required number of master mixes (one per siRNA), 250 nM siRNA was added, and the mixture transferred to Nucleocuvette™ Vessels (one per replicate). Electroporation was carried out in the 4D-Nucleofector™ X Unit, program EN-120, before the cells were diluted in RPMI and transferred to T75 culture flasks (Corning; one per replicate) and cultured as indicated.

### Cell viability assays

LNCaP or MR49F cells were transfected with siRNA overnight in 24-well plates at a density of 3x10^4^ or 4x10^4^ cells/well respectively, then cultured or treated as indicated for 3 or 6 days and counted as described previously^27^. Fatty acids, *cis*-vaccenic acid or oleic acid were supplemented as indicated for rescue studies. The concentration for supplementation was selected based on pilot cell viability experiments to determine the highest non-toxic dose of each fatty acid. For treatment only experiments LNCaP, MR49F, V16D, PNT1A or LOVO cells were seeded in 24-well plates at a density of 3x10^4^, 4x10^4^, 2.5x10^4^, or 2x10^4^ cells/well, respectively, then treated as indicated for 3 or 6 days and counted as described previously^27^.

### Colony formation

LNCaP cells were plated in 6-well plates (500 cells/well) in media supplemented with 100 µM *cis*-vaccenic acid (Cayman Chemical) and were incubated for two weeks at 37°C. Cells were then washed with PBS, fixed with 10% neutral buffered formalin and stained with 1% crystal violet for 30 mins. Colonies with > 50 cells were counted manually in a blinded manner.

### Ozone-induced dissociation (OzID) mass spectrometry

#### Cell lipid extraction

Cellular lipids for OzID-mass spectrometry and sum composition analyses were extracted from cells according to previously described literature^28^ and were quantified using deuterated lipid internal standards (SPLASH Lipid-o-mix®, Avanti Polar Lipids). In short, ∼5 million cells were cultured, transferred to 2 mL glass vials through trypsinisation, spun down and washed with PBS. To each sample vial, 220 µL methanol, 720 µL methyl-tert-butyl ether (MTBE; containing 0.1% butylated hydroxytoluene) and 40 µL SPLASH internal standard was added. Sample vials were capped and agitated for 1 h. 200 µL of aqueous ammonium acetate was added to each vial, and samples were vortexed before 5 min centrifugation at 5000 rpm. Sample supernatants were collected and stored at -20°C prior to mass spectrometric analyses.

#### Glycerophospholipid ozone-induced dissociation mass spectrometry

The sites of unsaturation in glycerophospholipids were elucidated using a high-resolution mass spectrometer (Orbitrap Elite, Thermo Scientific), modified to allow for ozone-induced dissociation (OzID) experiments. In line with previous literature^29^, ozone was produced via a generator (Titan-30UHC Absolute Ozone), which, through placement of a diverter valve on the nitrogen gas inlet to the higher-energy collisional dissociation (HCD) cell, replaced nitrogen gas to the cell with high concentration ozone gas (∼300 g/Nm^3^) in oxygen.

OzID mass spectrometry of lipids was conducted based on previously described methods^25^. All statistical comparisons were conducted using Microsoft Excel. Error bars report the standard error of the mean (SEM) at the 95th percentile confidence interval. Two-tailed student t-tests were calculated via conventional mathematical equations and are reported using standard asterisk nomenclature (*p<0.05, **p<0.01 and ***p<0.001).

### Patient derived explant tissue culture

Patient-derived explant (PDE) experiments were conducted according to Declaration of Helsinki principles, under protocols approved by the Human Research Ethics Committee of St Andrew’s Hospital (#80) and the University of Adelaide (H-2012-016). PDE culture was carried out using techniques established in our laboratory, as described previously^30^. Biopsy cores (6 mm/8 mm) were collected from men undergoing robotic radical prostatectomy at St. Andrew’s Hospital (Adelaide, SA) with written informed consent through the Australian Prostate Cancer Bio Resource. The tissue was dissected into smaller 1 mm^3^ pieces and cultured on Gelfoam sponges (Pfizer) in 24-well plates pre-soaked in 500 mL RPMI-1640 with 10% FBS, antibiotic/antimycotic solution. A939572 (10 µM) (Tocris Bioscience) was added into each well and the tissues were cultured in 5% CO_2_ in a humidified atmosphere at 37°C for 48h, then snap frozen in liquid nitrogen and stored at 80°C, or formalin-fixed and paraffin-embedded.

### Immunohistochemical quantification

Paraffin-embedded tissue sections (2–4 mm) were deparaffinized in xylene, rehydrated through graded ethanol, and blocked for endogenous peroxidase before being subjected to heat-induced epitope retrieval. IHC staining was performed using Ki67 and cleaved caspase 3 antibodies and the 3,3’-Diaminobenzidine (DAB) Enhanced Liquid Substrate System tetrahydrochloride (Merck Millipore, Darmstadt, Germany) as described previously^30^.

### Quantitative real-time PCR

RNA was extracted using the RNeasy Mini Kit (Qiagen) according to the manufacturer’s instructions, and concentration assessed using a Nanodrop DM-1000 spectrophotometer (NanoDrop Technologies). RNA was reverse transcribed using Superscript II RT and random hexamer primers according to the manufacturer’s instructions (BioRad). qRT-PCR was performed in triplicate with the BioRad C1000 Touch™ Thermal Cycler and CFX384™ Real-Time System using SYBR Green PCR Master Mix (BioRad) together with sequence specific primers, detailed in Supplementary Table S2. Expression levels were normalised against *IPO8* or *GAPDH* as reference genes.

### Western blotting

Protein lysates were collected in RIPA lysis buffer (10 mM Tris, 150 mM NaCl, 1 mM EDTA, 1% TritonX-100, 10% protease inhibitor). Western blotting on whole cell protein lysates were performed as previously described^27^. Protein levels were measured using primary antibodies detailed in Supplementary Table S1.

### Generation of ELOVL5 overexpressing cells (hELOVL5+)

LNCaP cells were transduced with the universal null control hRNA lentivirus [pLenti-TetCMV(hRNA(Null-Control))-Rsv(GFP-Puro)] or ELOVL5 hRNA lentivirus [pLenti-TetCMV(hRNA(ELOVL5))-Rsv(GFP-Puro)] designed by GenTarget Inc according to the manufacturer’s instructions. Briefly, 1-5x10^5^ cells were seeded in 24-well plates overnight and then treated with hexadimethrine bromide (8 µg/mL) and then transduced at MOI of 5. After 3 days, cell selection was undertaken using 4 µg/mL puromycin dihydrochloride (Gibco). ELOVL5 upregulation was confirmed by Western blot as described previously^27^.

### Oil Red O staining

LNCaP cells were treated with *cis*-vaccenic acid or oleic acid for 72 hr in 24-well plates. Cells were fixed in 10% neutral buffered formalin before being stained with 60% Oil Red O (Sigma-Aldrich) in water for 20 mins at room temperature in the dark. Cells were then washed with water and the Oil Red O extracted by isopropanol before absorbance was measured at 490 nm.

### Shotgun lipidomic analyses

Lipid extraction, mass spectrometry and data analysis for shotgun lipidomic analysis of samples were completed as described previously^31^.

#### Lipid extraction

Cells or tissues were homogenised in 800 µL ice-cold phosphate-buffered saline (PBS). Cell/tissue lysate (100 µL) was set aside to quantify DNA for normalisation before the remaining 700 µL was mixed with 800 µL 1 N HCl:CH_3_OH 1:8 (v/v), 900 µL CHCl_3_ and 200 mg/mL of the antioxidant 2,6-di-tert-butyl-4-methylphenol (BHT; Sigma Aldrich). 3 µL of SPLASH LIPIDOMIX Mass Spec Standard (#330707, Avanti Polar Lipids) was spiked into the extract mix. The organic fraction was evaporated using a Savant Speedvac spd111v (Thermo Fisher Scientific) at room temperature and the remaining lipid pellet was stored at -20°C under argon.

#### Mass spectrometry

Lipid pellets were reconstituted in 100% ethanol. Lipid species were analysed by liquid chromatography electrospray ionization tandem mass spectrometry (LC-ESI/MS/MS) on a Nexera X2 UHPLC system (Shimadzu) coupled with hybrid triple quadrupole/linear ion trap mass spectrometer (6500+ QTRAP system; SCIEX, Concord, Canada). Chromatographic separation was performed on a XBridge amide column (150 mm x 4.6 mm, 3.5 mm; Waters) maintained at 35 °C using mobile phase A [1 mM ammonium acetate in water-acetonitrile 5:95 (v/v)] and mobile phase B [1 mM ammonium acetate in water-acetonitrile 50:50 (v/v)] in the following gradient: (0–6 min: 0% B→6% B; 6–10 min: 6% B→25% B; 10–11 min: 25% B→98% B; 11–13 min: 98% B→100% B; 13–19 min: 100% B; 19–24 min: 0% B) at a flow rate of 0.7 mL/min which was increased to 1.5 mL/min from 13 min onwards. SM, CE, CER, DCER, HCER, LCER were measured in positive ion mode with a precursor scan of *m/z* 184.1, 369.4, 264.4, 266.4, 264.4 and 264.4 respectively. TG, DG and MG were measured in positive ion mode with a neutral loss scan for one of the fatty acyl moieties. PC, LPC, PE, LPE, PG, LPG, PI, LPI, PS and LPS were measured in negative ion mode by fatty acyl fragment ions. Lipid quantification was performed by scheduled multiple reactions monitoring (MRM), the transitions being based on the neutral losses or the typical product ions as described above. The instrument parameters were as follows: Curtain Gas = 35 psi; Collision Gas = 8 a.u. (medium); Ion Spray Voltage = 5500 V and 4,500 V; Temperature = 550 °C; Ion Source Gas 1 = 50 psi; Ion Source Gas 2 = 60 psi; Declustering Potential = 60 V and 80 V; Entrance Potential = 10 V and 10 V; Collision Cell Exit Potential = 15 V and 15 V. The following fatty acyl moieties were taken into account for the lipidomic analysis: 14:0, 14:1, 16:0, 16:1, 16:2, 18:0, 18:1, 18:2, 18:3, 20:0, 20:1, 20:2, 20:3, 20:4, 20:5, 22:0, 22:1, 22:2, 22:4, 22:5 and 22:6 except for TGs which considered: 16:0, 16:1, 18:0, 18:1, 18:2, 18:3, 20:3, 20:4, 20:5, 22:2, 22:3, 22:4, 22:5, 22:6.

*Data analysis:*

Peak integration was performed with the MultiQuant software version 3.0.3. Lipid species signals were corrected for isotopic contributions (calculated with Python Molmass 2019.1.1) and were normalized to internal standard signals. Unpaired t-test p-values and FDR corrected p-values (using the Benjamini/Hochberg procedure) were calculated in Python StatsModels version 0.10.1.

### ^13^C-fatty acid tracer experiments

#### Sample preparation and lipid extraction

LNCaP cells were transfected with siRNA in triplicate in 6-well plates at a density of 5x10^5^ cells/well for 4 days. 24 h post transfection, cells were supplemented with unlabelled or labelled media containing [^13^C_18_]-oleic acid, or [^13^C_16_]-palmitoleic acid as indicated. Following incubation with ^13^C-fatty acid containing medium for 72 h, media was aspirated and cells were washed with 4 °C aqueous sodium chloride (0.9% w/v NaCl/H_2_O), and scraped with 300 µL extraction buffer 1:1 LC-MS methanol:water (Optima®, Thermo Fisher Scientific) into a 1.5 mL microcentrifuge tube. Remaining cells were washed into the tube with an additional 300 µL extraction buffer, before 600 µL chloroform (Honeywell Research Chemicals) was added and the tubes vortexed. Samples were incubated on ice for 10 mins before being briefly vortexed and then centrifuged at 15,000 x *g* for 10 mins at 4°C. The lower aqueous layer was collected and dried under nitrogen flow for lipidomic LC-MS analysis.

#### Mass spectrometry

Dried lower phase was resuspended in 100 µL isopropanol:methanol:chloroform (4:2:1 v/v) buffer containing 7.5 mM ammonium formate. Samples were analysed on a Shimadzu Nexera-ZenoTOF 7600 LC-MS system (SCIEX, Concord, Canada) using an InfinityLab Poroshell 120 EC-C18 2.1 x 150 m 2.7 µm column. The mobile phase A was acetonitrile:water (6:4 v/v) and mobile phase B was isopropanol:acetonitrile (9:1 v/v), both of which contained 10 mM ammonium formate and 0.1%(v/v) formic acid. A 30-min gradient elution was carried out at 200 µL/min and in ambient temperature, starting with 25% B at 0 min, 25% B at 2 min, 60% B at 5 min, 90% B at 16 min, 90% B at 22 min, and 25% B at 23 min.

To quantify ^13^C-fatty acids incorporation into PG and CL, samples were analysed in negative mode at 4500 V with the MS1 mass range set to 700-1600. MS^2^ was acquired in IDA mode, in which top 20 ions after dynamic background subtraction were fragmented by CID with collision energy of 65 V and the zeno pulsing on.

To quantify double bond location and 13C enrichment of specific CL, PG and PC species, samples were analysed by MRM HR using EAD (EIEIO) fragmentation^32^ in positive mode using a spray voltage of 5500 V. EAD parameters were electron beam current 7000 nA, kinetic energy 11 keV, collision energy 15-40 V, reaction time 30 ms, accumulation time 0.3-0.4 s, and zeno pulsing on.

#### Data analysis

To confirm retention times of PG and CL species, instrument data files were converted into ABF format using Reifycs ABF Converter and analysed using MS-DIAL^33^. Then, to quantify ^13^C-enrichment and infer double bond location, the same files were converted into mzML format using Proteowizard’s MSConvert^34^ and analysed in MATLAB using in-house scripts.

### Flow cytometry studies

#### Mitochondrial superoxide detection

LNCaP or MR49F cells were transfected with siRNA for 4 days and supplemented with 10 µM *cis*-vaccenic acid for 24 h following transfection in 6-well plates. Cells were collected into fluorescence-activated sorting tubes and stained with 2.5 µM MitoSOX™ Red Mitochondrial Superoxide Indicator (Invitrogen) in PBS for 30 mins at 37 °C. Cells were centrifuged and washed with PBS three times before analysis using a BD FACSymphony™ A5 Cell Analyzer (BD Biosciences).

#### Apoptosis

LNCaP cells were transfected with siRNA for 4 days and treated with venetoclax 24 h following transfection in 24-well plates. Positive control cells were treated with 1 mg/mL doxorubicin hydrochloride 72 h before collection. Cells were collected into fluorescence-activated sorting tubes and co-stained with 5 µM 7AAD (Invitrogen) and 1X BD Pharmingen™ PE Annexin V (BD Biosciences) in FACS binding buffer for 20 mins on ice, before analysis using a BD LSRFortessa X20 flow cytometer (BD Biosciences).

### Mitochondrial isolation

LNCaP cells were nucleofected with siRNA for 4 days, before 1.4x10^7^ cells were harvested per group by trypsinisation and washed twice with ice-cold PBS. Mitochondrial isolation was carried out by differential centrifugation using the Mitochondrial Isolation Kit (MITOISO2; Sigma-Aldrich) according to the manufacturer’s instructions. Briefly, the required number of cells were resuspended to uniform suspension in Cell Lysis Buffer and incubated on ice for 5 mins (mixed at 1-minute intervals by pipetting up and down once). Two volumes of 1 x Extraction Buffer A were added, before samples were centrifuged at 600 x *g* for 10 mins at 4 °C. Supernatant was transferred to a new tube and centrifuged at 11,000 x *g* for 10 mins at 4 °C. Supernatant was collected as the cytosolic fraction, and the pellet was resuspended in a buffer suitable for application, as the mitochondrial fraction.

### Cytochrome c measurement

Isolated mitochondrial fractions were treated with 250 mg/mL L-ascorbic acid (Sigma-Aldrich) for 5 mins before cytochrome c content was determined by absorbance at 550 nm using a spectrophotometer.

### Electron microscopy

Electron microscopy fixing, embedding and imaging was carried out as previously described^6^. LNCaP cells transfected with siRNA were fixed after 72 h in 1.25% glutaraldehyde, 4% sucrose, 4% paraformaldehyde in PBS (pH 7.2) for 1 hour at room temperature, post-fixed in 1% osmium tetraoxide in water, dehydrated in a graded ethanol series, and embedded in epoxy resin Embed812 (Electron Microscopy Science). Ultrathin sections (70 nm) were cut on a Leica UCT6 ultramicrotome, collected on formvar-coated copper-iridium slot grids, counterstained with 2% uranylacetate and Reynold’s lead citrate, and imaged in an FEI Tecnai G2 Spirit transmission electron microscope at 80 kV.

### Confocal microscopy

LNCaP cells were transfected with siRNA in an 8-well Ibidi chamber (Ibidi, Fitchburg, WI, USA), at a density of 1.2x10^4^ cells/well, for 72 hours. Cells were stained with 1 µM MitoTracker™ Deep Red FM for 2 minutes before the stain was removed for subsequent staining with DMN-LD lipid droplet stain^35^ at 2 µM and immediate imaging. All fluorescence microscopy was performed on the Nikon A1+ confocal microscope (Nikon, Tokyo, Japan) equipped with a LU-N4/LU-N4S 4-laser unit (403, 488, 561 and 638 nm), using a Plan Apo λ 60× oil-immersion objective lens (1.4 aperture) at 1.2 AU pinhole with NIS Elements software (v4.5, Nikon). The A1-DUG GaAsP Multi Detector Unit employed to acquire images at 3× scanner zoom, at 512 px with z-stacks of 0.25 µm step depth. For each experiment, two wells were imaged per siRNA, with at least 13 cells captured per well. MitoTracker™ Deep Red FM signal was used to model the mitochondria using Imaris x64 (v9.5.1) “Surface function” with the following parameters. Surface Grain Size = 0.200 µm; Diameter of Largest Sphere = 0.250 µm; Enable Automatic Threshold = true; Classify Surfaces – “Number of Voxels Img = 1” above 10.0. DMN-LD signal was used to model lipid droplets using Imaris x64 (v9.5.1; Adelaide Microscopy, The University of Adelaide, Australia) “Spots function” with the following parameters. Estimated XY Diameter = 0.700 µm; Estimated Z Diameter = 1.40 µm; Classify Spots – “Intensity Max Ch=1 Img=1” above 1000. For the complete list of parameters for both functions see Supplementary Table S3. Distances between mitochondria and lipid droplets were calculated using from the “Surface function” using the “Shortest Distance to Spots” statistic. The percentage of mitochondria within 0.08 µm of a lipid droplet was reported for each micrograph, as membrane contact sites are generally defined with a 10-80 nm distance^36^).

### Statistical analysis

Statistical analysis was carried out using GraphPad Prism software v9.0.0 (2020, GraphPad Software), and data are representative of at least two independent experiments. Bar graphs and growth curves represent the mean ± SEM of at least two biological replicates unless indicated otherwise. For cell line studies, significance was measured by two-tailed unpaired *t*-test, one-way ANOVA or two-way ANOVA with multiple comparison tests as indicated. Significance is expressed as *, p<0.05; **, p<0.01; ***, p<0.001; ****, p<0.0001.

## Results

### The MUFA product of ELOVL5, cis-vaccenic acid, enhances prostate cancer cell viability

We have previously reported reduced cell viability in prostate cancer cells depleted of ELOVL5 that was rescued by *cis*-vaccenic acid (cVA), the only known monounsaturated fatty acid (MUFA) product of this enzyme^6^. We hypothesised that the pro-proliferative action of cVA (18:1*n*-7, *cis*) in ELOVL5-depleted prostate cancer cells was specific to this species of MUFA and could not be replicated by the closely related fatty acid oleic acid (OA; 18:1*n*-9, *cis*) (Fig. 1A). In two prostate cancer cell lines, hormone-sensitive LNCaP cells and treatment-resistant MR49F cells, cVA (10 µM) rescued the decrease in cell viability due to ELOVL5 depletion while OA (10 µM) did not (Fig. 1B). Increasing the OA concentration (30 µM) did not alter this effect (Fig. S1A), indicating that this effect of cVA was not simply due to the MUFA identity of cVA. Furthermore, exogenous cVA supplementation increased viability of both LNCaP and MR49F cells, but not the benign cell line PNT1A (Fig. 1C), with complimentary changes in cell death (Fig. S1B). This indicated a potentially selective importance of cVA to malignant prostate cells. Finally, cVA treatment caused enhanced colony formation in LNCaP cells (Fig. 1D), supporting a pro-proliferative role for this specific MUFA.

Monitoring cVA abundance in cells by mass spectrometry alone can be challenging due to the greater relative abundance of the OA isomer that shares an identical mass-to-charge ratio. Ozone-induced dissociation (OzID) mass spectrometry uses ozone to fragment carbon-carbon double bonds within lipids and can therefore be used to reveal changes in the abundance of individual structural isomers^25^. Using phosphatidylcholine (PC) lipids as proxies for MUFA isomer measurement, we confirmed that ELOVL5 knockdown reduced 18:1*n*-7, *cis* (cVA, ELOVL5 product) abundance in PC 34:1 (predominantly 16:0_18:1) and PC 36:1 (predominantly 18:0_18:1) (Fig. 1E; Fig. S1B), and increased 16:1*n*-7, *cis* (palmitoleic acid (POA), ELOVL5 substrate) abundance in PC 32:1 (predominantly 16:0_16:1) (Fig. 1E; Fig. S1C). These reported acyl isomers were dominant in each PC species, representing more than 85% of the isomeric acyl chains detected (Fig. S1C)

### The MUFA species produced by SCD1 are required for prostate cancer cell viability in vitro and ex vivo

Considering the general importance of MUFAs in cancer, and specifically the MUFA product of ELOVL5, cVA, in prostate cancer cells, we sought to further characterise the functional reliance of a range of prostate cancer models on cVA and OA production. The desaturase enzyme *SCD1* is often overexpressed in cancer and can promote the malignant phenotype^15^, and, accordingly, *SCD1* mRNA and protein is increased in our panel of prostate cancer cell lines, compared to PNT1A cells (Fig. S2A and B). Therefore, we used the small molecule inhibitor A939572 (Tocris Bioscience), to inhibit SCD1 activity^37^ in a range of prostate cancer cell lines (LNCaP, MR49F, V16D and 22Rv1). The efficacy of A939572 to suppress SCD1 activity was confirmed using shotgun lipidomics, whereby the ratio of SCD1 substrate to product (i.e. stearic acid-to-OA ratio) was increased in both LNCaP and MR49F cell lines (Fig. S2C). In addition, the OzID lipidomics workflow demonstrated reduced abundance of *n*-7 and *n*-9 MUFA isomers in PC species containing 16:1 or 18:1 acyl chains (Fig. 2A; Fig. S2D).

**Figure 2.**
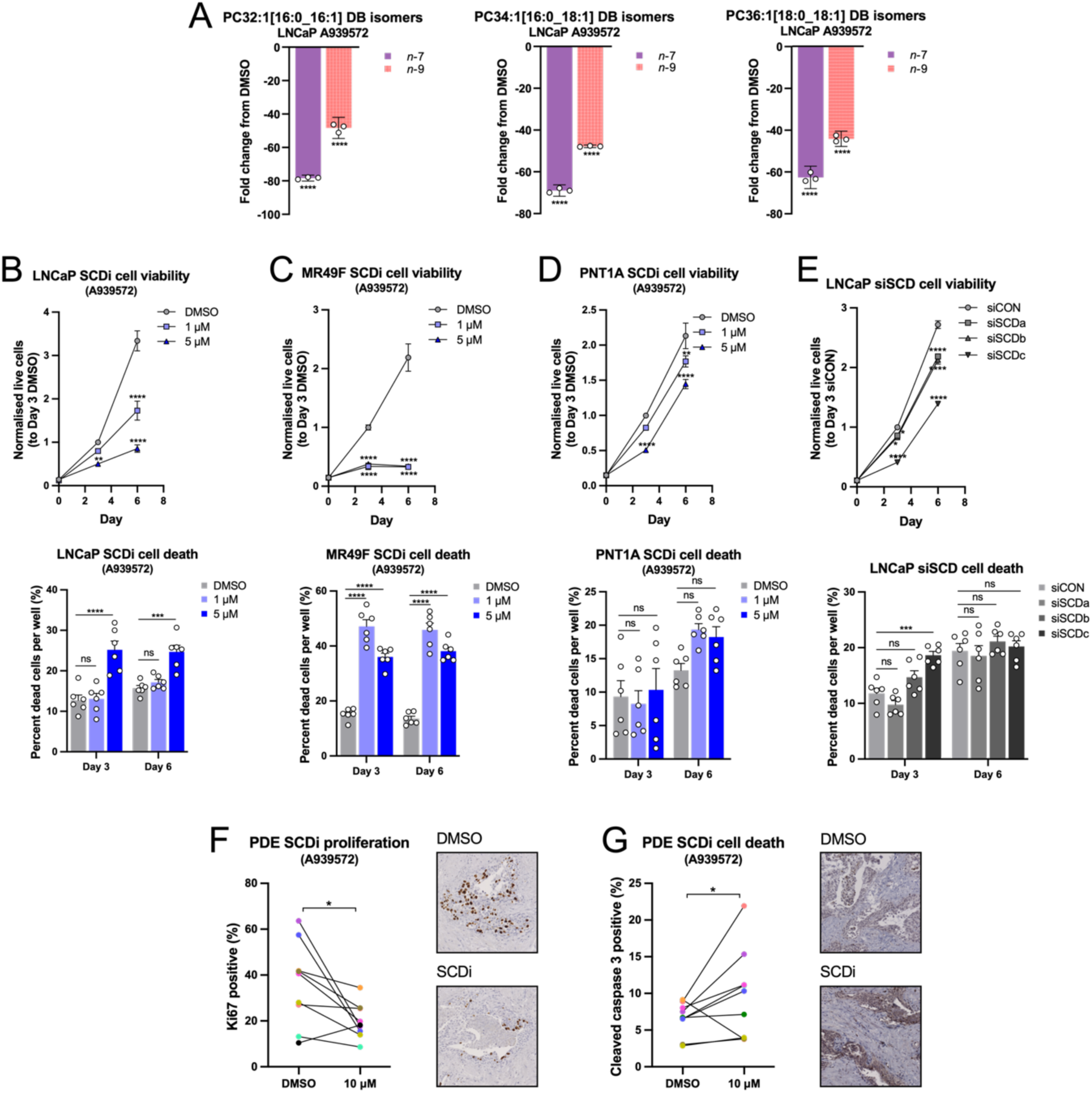
The MUFA species produced by SCD1 are required for prostate cancer cell viability *in vitro* and *ex vivo*. [A] Ozone-induced dissociation (OzID) lipidomic analysis of the relative *n*-7 and *n*-9 isomer abundance in phosphatidylcholines PC 32:1, PC 34:1 and PC 36:1 in A939572-treated LNCaP cells, 3 days post-treatment. Data are presented as mean ± 95% confidence interval (CI) (N = 1; n = 3). [B, C & D] Assessment of the effect of A939572 treatment on cell viability and cell death in prostate cancer cell lines LNCaP and MR49F, and benign prostate cell line PNT1A, three- and six-days post-treatment. Data are presented as normalised mean ± SEM (N = 2; n = 3). [E] Cell viability and cell death measurements under SCD1 knockdown (siSCD) in LNCaP prostate cancer cell line, three- and six-days post-treatment. Data are presented as normalised mean ± SEM (N = 2; n = 3). [F & G] Ki67 and cleaved caspase 3 immunostaining of patient-derived explant samples and quantification of the effect of A939572 treatment on proliferation [E] and cell death [F], 48 hours after treatment. Data are representative of nine independent patients (N = 9). Statistical significance was determined using two-way ANOVA and multiple comparisons [A – E], or student’s paired t-test [F & G]. * p<0.05; ** p<0.01; *** p<0.001; ****p<0.0001.

In all cell lines tested, SCD1 inhibition caused a dose-dependent reduction in cell viability and an increase in cell death (Fig. 2B and C; Fig. S2E and F). Inhibition of SCD1 in PNT1A cells reduced cell viability to a lesser extent than in malignant lines and did not increase cell death (Fig. 2D). Importantly, our observations made in LNCaP and PNT1A cell lines were confirmed using an independent SCD1 inhibitor, CAY10566^38^ (Cayman Chemical) (Fig. S2G and H), and three siRNAs against SCD1 (LNCaP cells only) (Fig. 2E).

The effects of SCD1 inhibition in cancer have been mainly addressed in cell line and animal models^15,39–41^, but not yet in the context of a clinical tumour microenvironment. To model clinical prostate cancer, we utilised our established patient-derived explant (PDE) culture approach, whereby tumour samples are biopsied following radical prostatectomy and cultured *ex vivo* over acute time points^42^. In PDEs (n = 9), SCD1 inhibition (A939572) reduced epithelial cell proliferation, by proliferative marker Ki67 immunostaining (Fig. 2F and S2I), and increased apoptosis, as determined by cleaved caspase 3 immunostaining (Fig. 2G and S2J).

### Cis-vaccenic acid can rescue the effects of SCD1 inhibition in prostate cancer cell lines

Having characterised the importance of MUFA production in prostate cancer cells and confirmed the reduction in cVA abundance with SCD1 inhibition, we sought to determine the contribution of cVA to regulation of these phenotypes. To achieve this, SCD1 inhibitor-treated cells were supplemented with cVA or OA. OA was used as a comparison, as the effects of SCD1 inhibition are usually rescued by this SCD1 product^39,40,43,44^. In both LNCaP and MR49F cell lines, however, supplementation of cVA could completely rescue the decrease in cell viability caused by SCD1 inhibition (Fig. 3A), while OA did not at equal concentration (10 µM). Increased OA concentration (30 µM) rescued the A939572-induced reductions in LNCaP and MR49F prostate cancer cell viability (Fig. S3A). Inhibition of SCD1 by CAY10566 was completely rescued by both cVA and OA (Fig. 3B), likewise for SCD1 knockdown (Fig. 3C). POA is the direct SCD1 product in the *n*-7 MUFA isomer family and is elongated by ELOVL5 to give cVA (Fig. 1A)^20,45^. For both SCD1 inhibitors, POA only partially rescued the decrease in cell viability (Fig. S3B), suggesting that SCD1 production of 18-carbon fatty acyl-CoA species is more important than 16-carbon species in prostate cancer cells.

**Figure 3.**
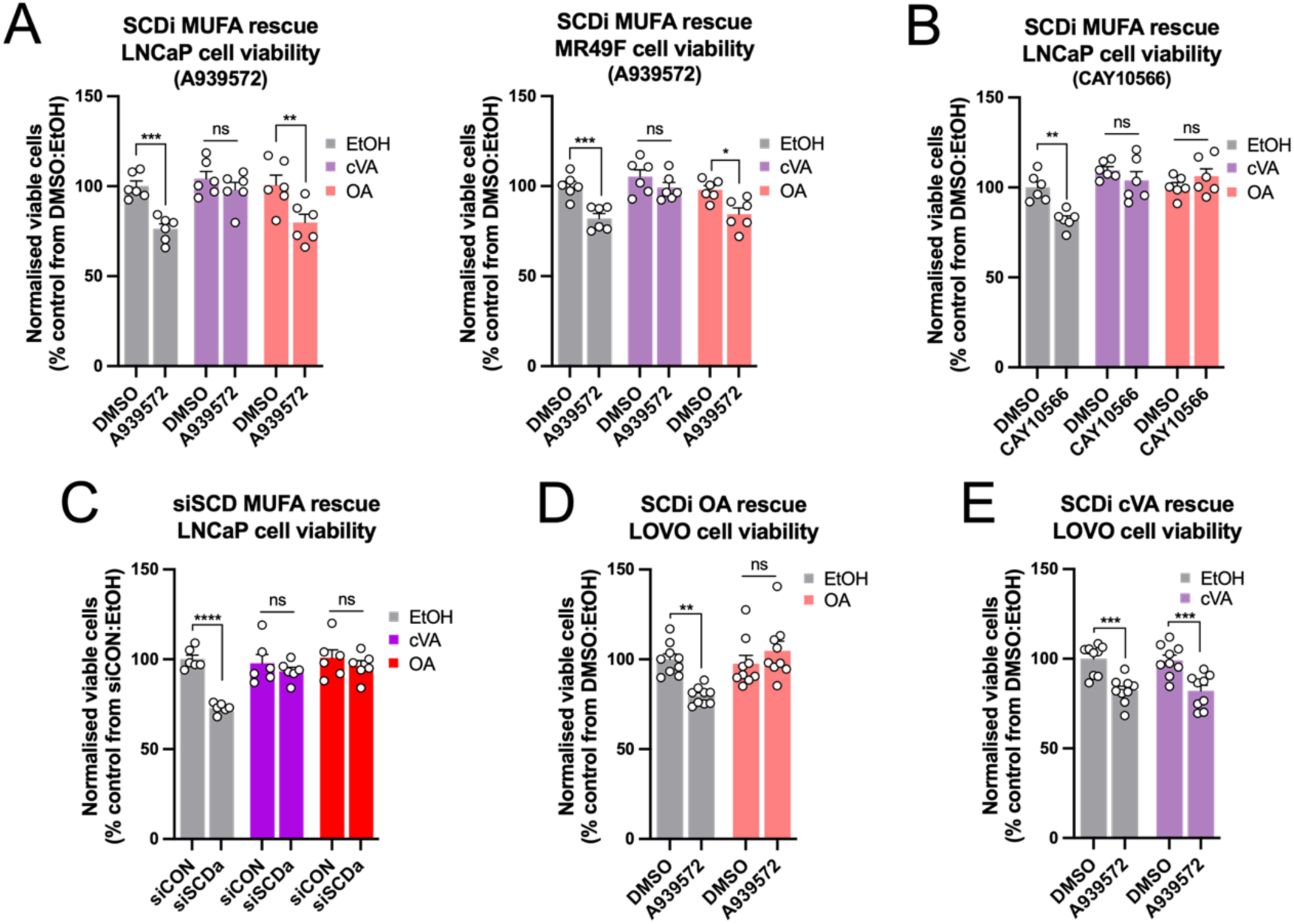
*Cis*-vaccenic acid and oleic acid can rescue the effects of SCD1 inhibition in prostate cancer cell lines. [A] Cell viability rescue experiments in LNCaP and MR49F cells for A939572 treatment-related changes in response to 10 µM *cis*-vaccenic acid (cVA) or oleic acid (OA), 96 hours post-transfection. Data are presented as normalised mean ± SEM (N = 2; n = 3). [B] Cell viability rescue experiments in LNCaP cells for CAY10566 treatment-related changes in response to 10 µM cVA or OA, 96 hours post-transfection. Data are presented as normalised mean ± SEM (N = 2; n = 3). [C] Cell viability rescue experiments in LNCaP cells for SCD1 knockdown-related changes in response to 30 µM cVA or OA, 96 hours post-transfection. Data are presented as normalised mean ± SEM (N = 2; n = 3). [D & E] Cell viability rescue experiments in LOVO colorectal cancer cell line for A939572 treatment-related changes in response to 10 µM OA [D] or cVA [E], 96 hours post-transfection. Data are presented as normalised mean ± SEM (N = 3; n = 3). Statistical significance for was determined using two-way ANOVA and multiple comparisons. * p<0.05; ** p<0.01; *** p<0.001; ****p<0.0001.

Next, we tested the sensitivity of *ELOVL5* overexpressing cells (hELOVL5+) to SCD1 inhibition, where increased abundance of ELOVL5 should result in larger cellular pools of cVA^20^, over acute time points. hELOVL5+ cells were more resistant to SCD1 inhibition (Fig. S3C), indicating that increased *ELOVL5* expression and cVA production can maintain prostate cancer cell viability under MUFA-deficient conditions.

The ability of cVA to rescue SCD1 inhibition-induced reductions in prostate cancer cell viability in any capacity was unexpected. As such, we aimed to recapitulate previously published data from a colorectal cancer cell line (LOVO), where the effects of A939572 was rescued by OA supplementation^46^. As expected, OA rescued the decrease in cell viability induced by A939572 in LOVO cells (Fig. 3D); however, cVA did not rescue the effects of A939572 in this context (Fig. 3E). This demonstrates the possibility of a tumour type-selective dependency on cVA. Interestingly, the ability of cVA to rescue SCD inhibition by A939572 correlated with the expression of *ELOVL5* compared to *ELOVL6* in LNCaP and LOVO cell lines (Fig. S3D).

To rule out differences in MUFA uptake, we assessed the ability of LNCaP and MR49F cells to take up cVA and OA. Oil Red O staining revealed that supplementation of culture media with low and high doses of cVA and OA caused the same increase in overall lipid content in both cell lines (Fig. S3E). In addition, 18:1 abundance increased in phospholipid species, triglycerides and sphingomyelins following cVA or OA supplementation in LNCaP cells (Fig. S3F and G), supporting equivalent uptake of each MUFA.

### Cis-vaccenic acid is incorporated into cardiolipins and cardiolipin precursors in prostate cancer cells

The role of cVA in cancer has not been determined, but there is some evidence that points to a unique enrichment of this unusual fatty acid in the mitochondrial-specific phospholipid, cardiolipin (CL)^25,47^. The biosynthesis of CLs (Figure 4A) requires the esterification of up to four acyl chains. Early evidence in rats suggested that among the acyl chains bound to CLs, 18:1*n*-7, cis (cVA) was the predominant 18-carbon MUFA isomer rather than the 18:1*n*-9, cis isomer (OA)^47^. This was supported by a recent report by Young and colleagues^25^ that the *n*-7 isomer is the most common 18:1 species in the CL precursor lipid phosphatidylglycerol (PG) in LNCaP prostate cancer cells^25^. Similarly, data from our cell line panel showed that the *n*-7 isomer was overrepresented in PG 34:1 (predominantly 16:0_18:1; Fig. 4B), compared to 34:1 species in other phospholipid classes (Fig. 4C, S4A and B). Treatment with A939572 decreased PG 34:1 abundance, and 18:1 acyl chain abundance across the entire PG class in LNCaP cells, and cVA supplementation partially rescued this decrease, while OA did not (Fig. 4D and E). In addition, A939572 treatment increased the abundance of saturated PG species and decreased monounsaturated PG abundance, while supplementation of cVA, but not OA, reversed these changes (Fig. 4F). PG is a precursor for mature CLs (Fig. 4A)^48^, and thus the unique ability of cVA to rescue changes in PG lipids raises the possibility of a selective role for cVA in regulating CL content in prostate cancer cells.

**Figure 4.**
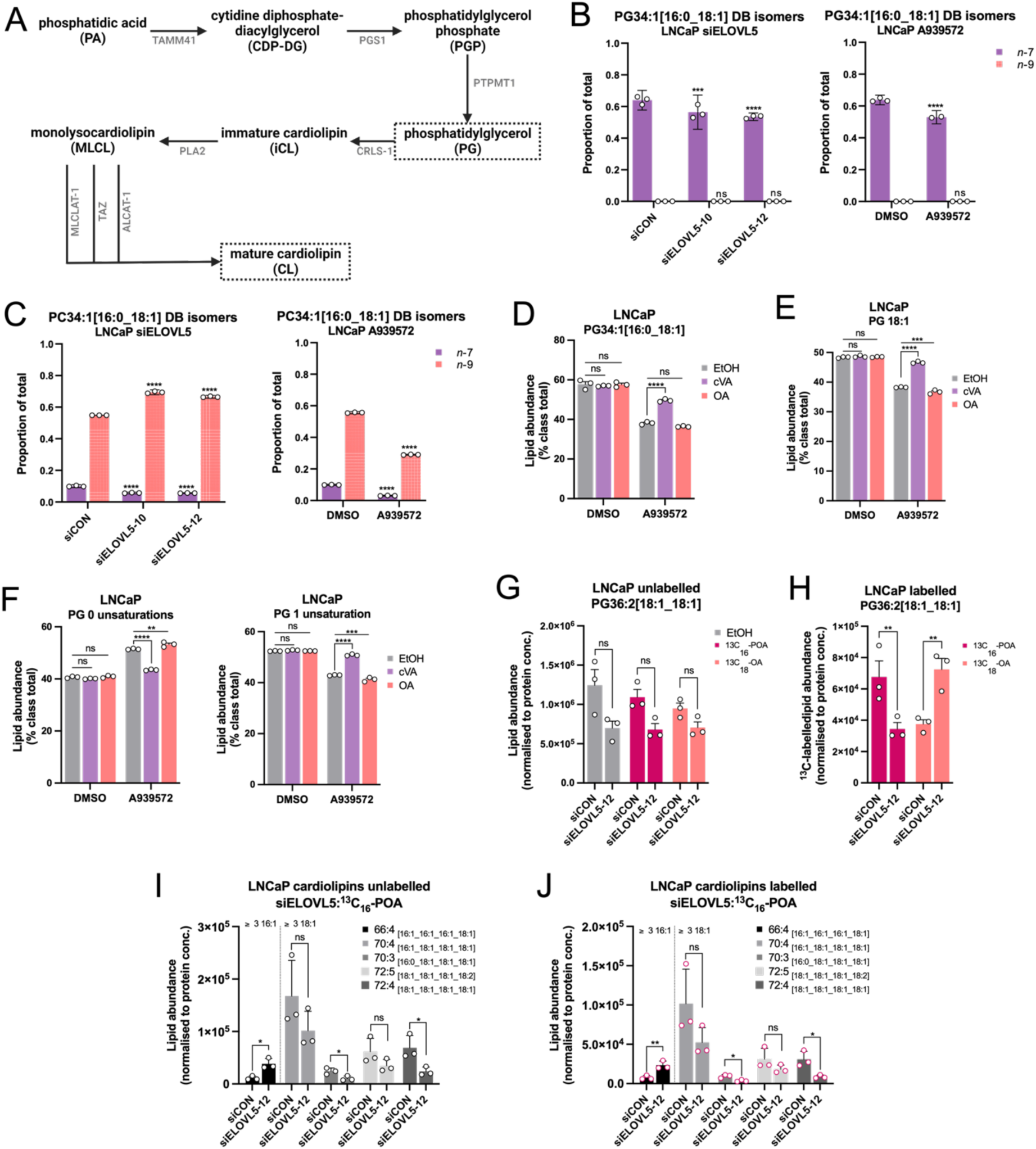
*Cis*-vaccenic acid is incorporated into cardiolipins and cardiolipin precursors in prostate cancer cells. [A] Cardiolipin (CL) synthesis and remodelling pathway diagram, highlighting the phosphatidylglycerol (PG) intermediate and CL product. ALCAT-1, acyl-CoA:lysocardiolipin acyltransferase 1; CRLS1-1, cardiolipin synthase 1; PGS1, phosphatidylglycerophosphate synthase 1; PLA2, phospholipase A2; PTPMT1, protein tyrosine phosphatase localized to the mitochondrion 1; MLCLAT-1, monolysocardiolipin acyltransferase 1; TAMM41, TAM41 mitochondrial translocator assembly and maintenance homolog; TAZ, tafazzin. [B & C] Ozone-induced dissociation (OzID) lipidomic analysis of relative *n*-7 and *n*-9 isomer abundance in phosphatidylglycerol (PG) 34:1 [B] and phosphatidylcholine (PC) 34:1 [C] in ELOVL5 knockdown (siELOVL5) and A939572-treated LNCaP cells. Data are presented as mean ± 95% confidence interval (CI) (N = 1; n = 3). [D – F] Lipidomic analysis of phosphatidylglycerol (PG) acyl chains in response to A939572 treatment in combination with 10 µM *cis*-vaccenic acid (cVA) or oleic acid (OA). Data are presented as mean ± SEM (N = 1; n = 3). [G] Lipidomic analysis of PG 36:2 abundance in LNCaP ELOVL5 knockdown cells. Data are presented as mean ± SEM (N=1; n=3). [H] Lipidomic measurement of ^13^C_16_-palmitoleic acid (POA) and ^13^C_18_-OA incorporation into PG 36:2 in LNCaP ELOVL5 knockdown cells. Data are presented as mean ± SEM (N = 1; n = 3). [I] Lipidomic analysis of CL abundance in LNCaP ELOVL5 knockdown cells. Data are presented as mean ± SEM (N=1; n=3). [J] Lipidomic measurement of ^13^C_16_-POA incorporation into CL species in LNCaP ELOVL5 knockdown cells. Data are presented as mean ± SEM (N = 1; n = 3). Statistical significance was determined by two-way ANOVA [B – J]. * p<0.05; ** p<0.01; *** p<0.001; ****p<0.0001.

To more directly investigate the contribution of cVA to PG and CL species, we carried out ^13^C-labelled fatty acid tracer lipidomics. Labelled cVA was not commercially available, so labelled POA (^13^C_16_-POA), the ELOVL5 substrate and direct precursor of cVA (Fig. 1A), was used to measure incorporation of this labelled fatty acid species into PGs and CLs as a substitute. The abundance of endogenous PG 36:2 (predominantly 18:1_18:1) was reduced by ELOVL5 knockdown (Fig. 4G) irrespective of the labelled fatty acid supplemented. Monitoring of PGs revealed a heavy labelled PG 36:2 post ^13^C_16_-POA supplementation (Fig. 4H), inferring that the labelled supplement was both elongated and incorporated into the PG 18:1_18:1 lipid species. Moreover, ELOVL5 knockdown reduced the enrichment into PG 36:2 (Fig. 4H). Independent supplementation of labelled OA (^13^C_18_-OA) also revealed incorporation into PG 36:2 but relative to ^13^C_16_-POA labelled PG 36:2 was less abundant and instead increased with ELOVL5 knockdown (Fig. 4H), possibly as an adaptation to overcome the effects of ELOVL5 targeting. In CLs, ELOVL5 knockdown increased the abundance of endogenous species containing more than three 16:1 acyl chains (e.g. CL 66:4) and trended towards decreased abundance for those containing more than three 18:1 acyl chains (e.g. CL 70:4, 70:3, 72:5 and 72:4) (Fig. 4I). Extracellular ^13^C_16_-POA was present in these specific CLs (Fig. 4I), and the enrichment was increased in 16:1-containing CLs and decreased in 18:1-containing CL species after ELOVL5 knockdown (Fig. 4J). Acyl chain identification confirmed the incorporation of ^13^C_16_-POA into the acyl chains of PG 36:2, and the CL species of interest (Fig. S4C). Detection of labelled ^13^C_16_-POA in 16:1 acyl chains indicated direct incorporation, while ^13^C_16_-POA in 18:1 acyl chains indicated an elongation event, providing evidence for ELOVL5 activity in PG and CL synthesis. Combined, these results demonstrate that cVA is a substrate for CL synthesis.

### Cis-vaccenic acid-containing cardiolipins regulate mitochondrial homeostasis in prostate cancer cell lines

CLs are found in the outer and inner mitochondrial membrane of the mitochondria and bind cytochrome c at the inner mitochondrial membrane^49,50^. CL content in the mitochondrial membrane is an important regulator of intrinsic apoptosis that mediates changes to the biophysical properties of the outer mitochondrial membrane, promoting mitochondrial outer membrane permeabilization (MOMP) and caspase activation^51,52^. In addition, CL are oxidised during cytochrome c release and initiation of apoptotic cell death signalling pathways^53,54^ (Fig. 5A). Analysis of cytochrome c abundance in mitochondria from the LNCaP prostate cancer cell line demonstrated no significant change with ELOVL5 knockdown (Fig. 5B and S5A). In addition, ELOVL5 knockdown did not sensitise LNCaP cells to a BH3 mimetic, venetoclax (Fig. 5C and S5B), indicating that altered CL cVA content did not induce changes in cytochrome c-related mitochondrial phenotypes in this prostate cancer cell line model and that the changes observed in cell viability with *ELOVL5* knockdown are likely independent of apoptosis, consistent with our previous observations^6^.

**Figure 5.**
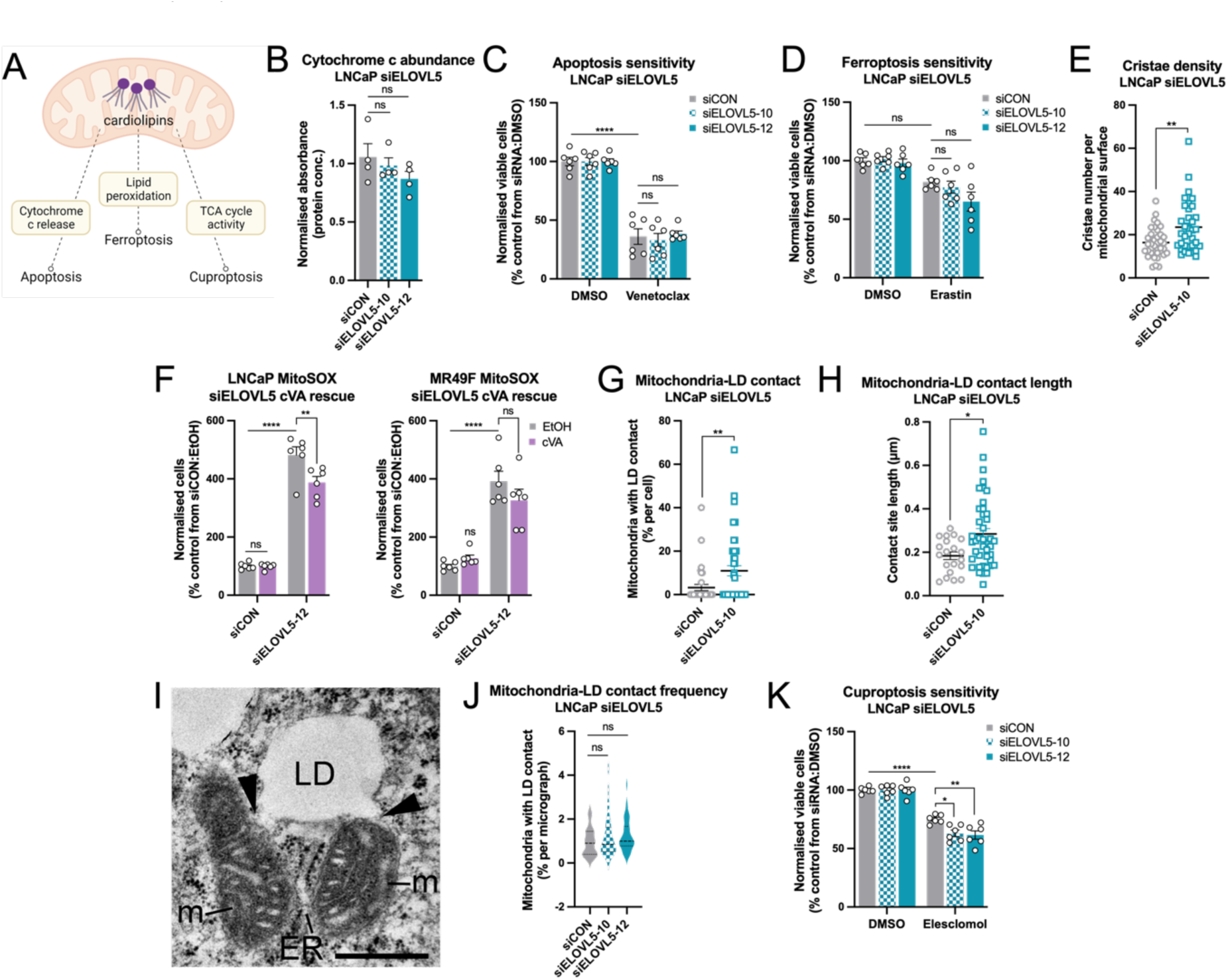
*Cis*-vaccenic acid-containing cardiolipins regulate mitochondrial homeostasis in prostate cancer cell lines. [A] Diagram depicting three programmed cell death pathways, apoptosis, ferroptosis and cuproptosis, that cardiolipins are linked to via three distinct mechanisms. [B] Cytochrome c abundance in ELOVL5 knockdown LNCaP cells measured by absorbance at 550 nm. Data are presented as mean ± SEM (N = 1; n = 2). [C] Cell viability of LNCaP ELOVL5 knockdown cells in response to 10 µM venetoclax treatment. Data are presented for each siRNA as normalised to DMSO, mean ± SEM (N = 2; n = 3). [D] Cell viability of LNCaP ELOVL5 knockdown cells in response to 10 µM erastin treatment. Data are presented for each siRNA as normalised to DMSO, mean ± SEM (N = 2; n = 3). [E] Electron microscopy analysis of mitochondrial cristae density changes in response to ELOVL5 knockdown in LNCaP cells. Data are presented as mean ± SEM (N = 1; n = 35). [F] Mitochondrial superoxide (MitoSOX) measurement for LNCaP and MR49F cells for ELOVL5 knockdown-related changes in response to 10 µM cVA. Data are presented as normalised mean ± SEM (N = 2; n = 3). [G & H] Electron microscopy analysis of mitochondria-lipid droplet (LD) contact frequency [G] and length [H] changes in response to ELOVL5 knockdown in LNCaP cells. Data are presented as mean ± SEM of one independent experiment (N = 1), with 35 and 45 (n = 35 and 45) [G], 20 and 42 (n = 20 and n = 42) [H] technical replicates for siCON and siELOVL5-10, respectively. [I] Representative electron microscopy image showing mitochondria-LD contacts in ELOVL5 knockdown cells. The scale bar is representative of 500 nm. [J] Fluorescence confocal microscopy quantitation of the percentage of mitochondria per micrograph with LDs less than 0.08 nm in response to *ELOVL5* knockdown in LNCaP cells, calculated using the “spots to surfaces” function on Imaris. [K] Cell viability of LNCaP ELOVL5 knockdown cells in response to 1 nM elesclomol treatment. Data are presented for each siRNA as normalised to DMSO, mean ± SEM (N = 2; n = 3). Statistical significance was determined by one-way ANOVA [B & J], two-way ANOVA [C, D, F & L], Mann-Whitney test [E] or student’s unpaired t-test [G & H]. * p<0.05; ** p<0.01; *** p<0.001; ****p<0.0001.

Another cell death pathway linked to altered phospholipid saturation profiles is ferroptosis, an iron-dependent form of cell death^55^. Ferroptosis can be initiated by the accumulation of lipid peroxide species, whereby PUFA-containing phospholipids are susceptible to peroxidation, due to the increased frequency of double bonds^3^. Moreover, lipid ROS accumulation specifically in mitochondria following ferroptosis induction has been reported previously^56^. Therefore, we utilised ferroptosis inducer erastin^55^ to investigate the effect of ELOVL5 knockdown, and altered CL content, on ferroptosis sensitivity. In LNCaP cells, erastin treatment reduced cell viability but did not increase cell death (Fig. 5D and S5C). Knockdown of ELOVL5 did not alter these effects of erastin (Fig. 5D and S5C), suggesting that the changes in CL content did not influence ferroptosis sensitivity.

We have previously reported that ELOVL5 and mitochondrial respiration, whereby ELOVL5 protein levels were positively correlated with mitochondrial oxygen consumption^6^. It is well established that dysregulation of CL homeostasis impacts mitochondrial cristae organisation, and thereby electron transport chain (ETC) activity^57^ and complex stabilisation^58^, and increased mitochondrial superoxide^59,60^. Here we show that ELOVL5 knockdown in LNCaP cells had increased cristae number per mitochondrial surface determined by electron microscopy (Fig. 5E). Next, we assessed whether this ELOVL5 loss-of-function induced alteration in morphology impacted mitochondrial superoxide production. ELOVL5 knockdown significantly increased mitochondrial superoxide production, and this was partially rescued by cVA supplementation (Fig. 5F). Moreover, electron microscopy analysis of mitochondria-organelle contacts demonstrated increased frequency (Fig. 5G) and length (Fig. 5H) of mitochondria-lipid droplet contacts in ELOVL5 knockdown LNCaP cells (Fig. 5I), which was confirmed using fluorescent confocal microscopy (Fig. 5J). This change in mitochondria-lipid droplet arrangement have been reported to be indicative of mitochondrial energy generation-related stress^61–63^.

These links between ELOVL5, CLs and mitochondrial energy production and morphology, prompted us to investigate sensitivity of these cells to an additional form of cell death, cuproptosis. Cuproptosis is a recently described a copper-dependent form of cell death^64^, potentiated by copper-induced aggregation of lipoylated proteins. Of relevance to this study, the few enzymes that are known to be lipoylated are all linked to the tricarboxylic acid (TCA) cycle initiation^65^, which produces co-factors required for ETC complex activity^66^. Elesclomol is a copper ionophore that induces cuproptosis by importing copper into the cell^64^. Interestingly, treatment of LNCaP cells with elesclomol reduced cell viability, and this was enhanced under ELOVL5 knockdown, after normalising for any effects of the knockdown alone (Fig. 5K). This was also associated with an increase in cell death (Fig. S5D). Overall, these findings suggest that mitochondria are sensitive to changes in ELOVL5 activity in prostate cancer cells.

### “ELOVL5 activity” in tumour tissue is associated with clinical outcome in patients with localised prostate cancer

To investigate the clinical relevance of our *in vitro* and *ex vivo* findings, we carried out lipidomic analysis on radical prostatectomy samples from patients with localised prostate cancer for whom the clinical outcome was known. A total of 56 patient tumour samples were analysed, with 41 experiencing no disease relapse (NR) over a follow-up period of up to 12 years, while the remaining 15 experienced clinical relapse (CR) defined as documented metastatic disease detected by conventional imaging (Table 1 and Fig. 6A). Total lipid abundance was significantly increased in the CR group compared to the NR (Fig. 6B). To crudely investigate cVA-related effects in these samples, we utilised the ratio of 18:1 to 16:1 acyl chains in various lipid species, to represent an “ELOVL5 elongation ratio”. We first calculated the “ELOVL5 elongation ratio” in all lipid species, and determined that there was a small but not significant increase in CR tumour tissue compared to NR (Fig. 6C). We then focussed on the abundant phospholipid species, namely phosphatidylcholine (PC), phosphatidylethanolamines (PE) and other species of interest, lysophosphatidylcholines (LPC) and 1-alkenyl,2-acylphosphatidylethanolamines (PE-P), and demonstrated that the “ELOVL5 elongation ratio” was increased in CR tumours, compared to NR tumours (Fig. 6D and E). Taken together, these clinical data suggests that increased lipid abundance and enhanced SCD1/ELOVL5 activity are associated with clinical outcome in patients with localised prostate cancer, supporting our *in vitro* results.

**Figure 6.**
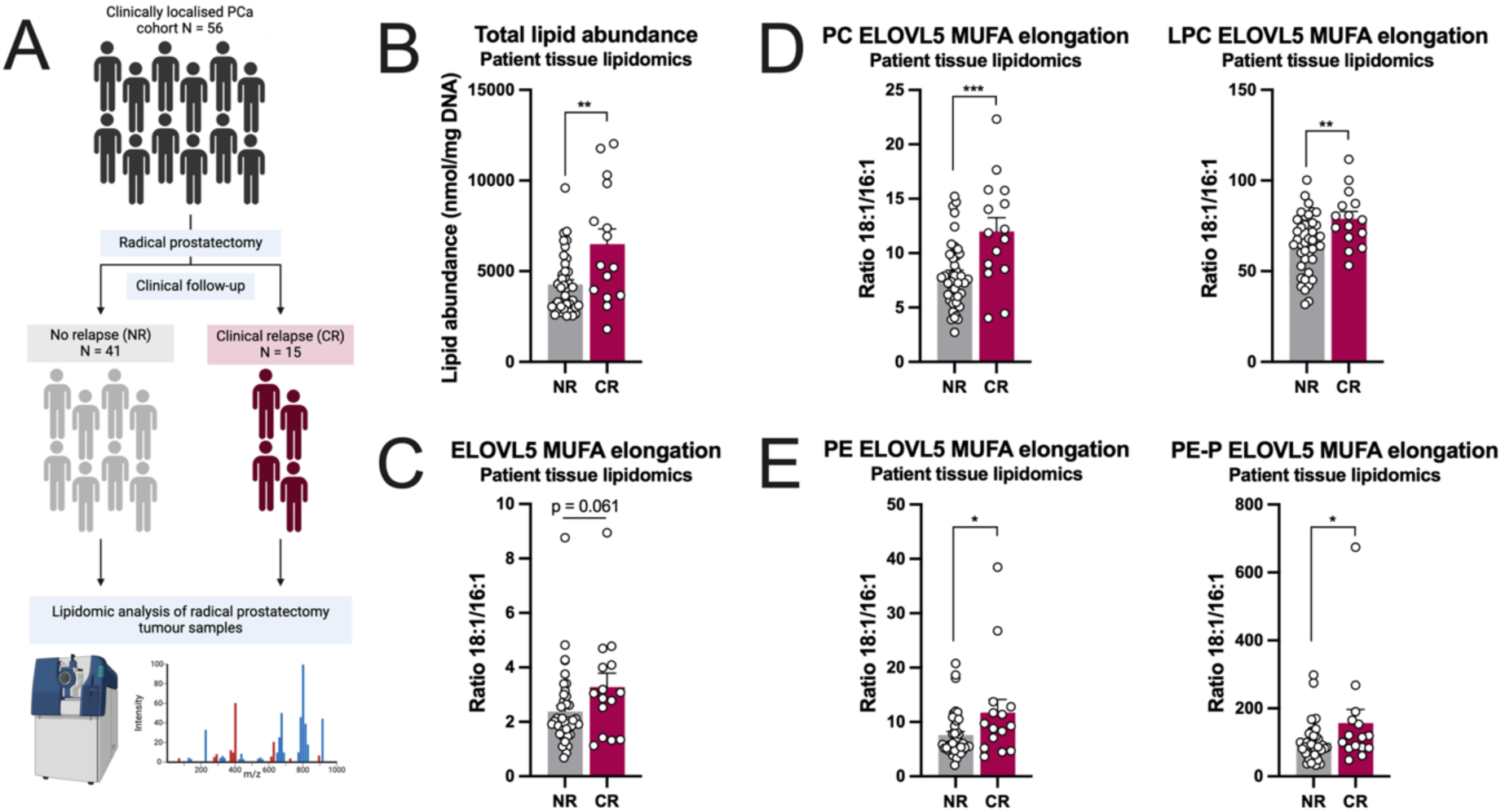
“ELOVL5 activity” in tumour tissue is associated with clinical outcome in patients with localised prostate cancer. [A] Men with clinically localised prostate cancer (N = 56) underwent radical prostatectomy, without prior treatment, at the Royal Adelaide Hospital or St Andrew’s Hospital (Adelaide, SA, Australia) and were followed-up to determine relapse status. No-relapse (NR) patients (N = 41) were followed up after a minimum of five years and did not experience biochemical (increased prostate specific antigen levels in the blood) or clinical relapse (metastatic disease). Clinical relapse (CR) patients (N = 15) were followed up and experienced biochemical relapse. They were then treated with a range of modalities, but ultimately developed clinical metastases that was occasionally terminal. Lipidomic analysis was carried out on patient tissue collected at radical prostatectomy. [B] Total lipid abundance in NR and CR tissue. [C – E] The ratio of 18:1-to-16:1 acyl chains (“ELOVL5 MUFA elongation ratio”) across all lipid species [C], in phosphatidylcholines (PC) and lysophosphatidylcholines (LPC) [D] and phosphatidylethanolamines (PE) and 1-alkenyl,2-acylphosphatidylethanolamines (PE-P) [E] in NR and CR tissues. in phosphatidylcholines (PC) in NR and CR tissues. Data are presented for B – E as mean ± SEM of 68 (N = 41) and 26 (N = 15) patients in the NR and CR groups, respectively. Statistical significance was determined by student’s unpaired *t*-test [B – D]. * p<0.05; ** p<0.01; *** p<0.001; ****p<0.0001.

**Table 1.** Clinical characteristics of the localised prostate cancer cohort.

|  | Median [first quartile, second quartile] or number |  |
| --- | --- | --- |
|  | Non-relapse (NR) | Clinical relapse (CR) |
| Number of men | 41 | 15 |
| Age, years | 60.4 [57.5, 65.3] | 65.1 [62.7, 67.8] |
| Gleason grade at surgery |  |  |
| ≤6 | 12 | 1 |
| 7 | 28 | 11 |
| 8 | 1 | - |
| ≥9 | - | 3 |
| PSA at surgery, ng/mL | 6.47 [5.00, 9.00] | 9.10 [7.50, 11.00] |
| Time of follow-up, months | 114.3 [98.6, 125.3] | 115.5 [90.6, 140.0] |

## Discussion

The importance of SCD1 expression and MUFA production in cancer cells is well-known^3,15^, however, the relative contributions of individual MUFA isomers to cancer progression is only recently being appreciated^25^. Here we show that an understudied 18:1 MUFA isomer, *cis*-vaccenic acid, is critical for prostate cancer cell viability and that it can be used to rescue the effects of specific cVA reduction and broad MUFA depletion in this setting. Historically, OA rescue of SCD1 inhibition has been demonstrated in many cancer types^16,39,40,67^, including prostate cancer^41^. Our observations indicate that other MUFA products of SCD1 are also important. While we are not the first to report on cVA in cancer^39,68–70^, to our knowledge this is the first study to conduct an in-depth mechanistic analysis of the role of cVA, beyond observing alterations to cVA abundance. Given that cVA is a direct product of ELOVL5^20^, and there is a unique importance of ELOVL5-mediated fatty acid elongation in prostate cancer^6^, there may be an increased relevance of cVA in prostate cancer, over other cancer types. Indeed, previous research has shown that androgen receptors (ARs) play a crucial role in prostate cancer progression and regulate key steps of *de novo* fatty acid synthesis, as well as fatty acid elongation and desaturation, including transcriptional regulation of *ELOVL5* and *SCD1* by AR^5,6^. While cVA may be simply a by-product of these effects of AR signalling that is being opportunistically utilised in malignant cells, the findings presented in this study suggest that it is more likely a consequence of aberrant ELOVL5 activity that confers a growth advantage to prostate cancer cells, over their benign counterparts. Specifically, the ability of cVA to increase cell viability and reduce cell death in our prostate cancer cell lines, but not the benign PNT1A cell line, is in support of this hypothesis.

The first report on the effect of cVA in cancer demonstrated a small reduction in cell viability of HT-29 colorectal cancer cell line with cVA treatment, at doses comparable to those used in this study^68^. Subsequent studies reported that red blood cell fatty acid content is a potential biomarker for cancer risk and showed a positive association between red blood cell *cis*-vaccenic acid (18:1*n*-7) and breast cancer risk in a Chinese population^69^ and a similar trend across a range of cancer types in a Spanish population^70^. Moreover, depletion of retinoblastoma protein in mouse embryonic fibroblasts induced widespread lipidomic changes, including increased expression of SCD1 and associated increased abundance of 18:1 fatty acids, including cVA^71^. While that investigation was not strictly carried out in a cancer model, inactivation of retinoblastoma pathway components is observed in almost all human malignancies^72^. Finally, a study conducted in a lung cancer cell line demonstrated that treatment with 100 µM cVA, along with other *cis*-MUFA species, rescued the effects of SCD1 inhibition on cell viability^39^, however the mechanism of this effect was not addressed. Therefore, despite these previous studies on cVA in cancer, the data presented in our study contributes novel insights into the pro-oncogenic role of cVA in prostate cancer^39^, including possible roles in cuproptosis and mediation of mitochondrial function.

The ability of SCD1-produced MUFA isomers to rescue SCD1 inhibition is likely to depend on several factors. For example, fatty acid elongase enzymes (ELOVLs) display distinct substrate specificity^45,73^, and hence differential expression of the seven *ELOVL* isoforms likely influence MUFA isomer production in different cancer types via regulation of SCD1 substrate availability. Of relevance to this study, prostate and colon tissues display differential expression of relevant *ELOVL* enzymes that correlate with the effects of cVA and oleic acid in each of these cancer types. Prostate tissue expresses high levels of *ELOVL5* relative to *ELOVL6*^74^, which is maintained in prostate cancer cells^6^. Conversely, colon tissue expresses more *ELOVL6* relative to prostate tissue^74^ and *ELOVL6* is overexpressed colon cancer compared to matched benign regions^75^. These patterns may favour cVA production in prostate cancer cells, and OA production in colorectal cancer cells (Fig. 1A) by inducing a reliance on each 18:1 MUFA species and facilitating the rescue of the effects of SCD1 inhibition in the respective model. In support of this hypothesis is our finding that the expression of *ELOVL5* relative to *ELOVL6* is much higher in LNCaP prostate cancer cells compared to LOVO colorectal cancer cells. Complementary to fatty acid elongase enzymes, expression of fatty acid desaturase enzymes, fatty acid desaturase 1 and 2 (FADS1 and FADS2) could also influence MUFA isomer preference between different cell types. Whilst generally attributed to the production of polyunsaturated fatty acids^76^, FADS1 and FADS2 can also be involved in MUFA production^25^. Noncanonical MUFA isomers formed by the apocryphal activity of FADS2 have recently been identified in cancer cells and appear to be protective against the effects of SCD1 inhibition^25,77^, since resistance to SCD1 inhibition correlates with high FADS2 expression^77^. Finally, the importance of ELOVL5-mediated production of cVA in prostate cancer cells may also be explained by the expression of *FASN*, and the transcriptional regulation of this enzyme by AR signalling^78^. The palmitic acid produced by FASN activity is lipotoxic to mammalian cells at high concentrations^79^. Consequently, SCD1 activity on palmitic acid may be increased in prostate cancer cells, to convert this potentially toxic SFA to POA. In addition, POA is a bioactive lipokine^23^, and as such prostate cancer cells may favour elongation to cVA to diminish any potential effects of POA signalling.

The data presented in this study presents cVA as a critical growth-promoting factor in prostate cancer, but did not consider the stereoisomer of cVA, *trans*-vaccenic acid (tVA), as this is a distinct dietary-derived fatty acid species, not modified by ELOVL5^80^. Dietary tVA was recently found to mediate anti-tumour effects via reprogramming of CD8+ T-cells^81^, suggesting that both forms of this understudied fatty acid are of interest and should be studied in the context of cancer. In contrast to tVA, our study identifies a unique role for cVA within prostate cancer cells themselves, as a key MUFA species in a mitochondrial specific phospholipid, CL. CLs are localised to both the outer and inner mitochondrial membranes (OMM and IMM, respectively)^82^, where their unique tetra-acyl conformation facilitates efficient membrane folding and formation of tightly packed cristae^57^, required for mitochondrial ATP production via the ETC^83^. While CLs are predominantly made up of 18:2 acyl chains, 18:1 acyl chains are prominent^84^, and have previously been identified as *n*-7 isomers in rat cardiac tissue^47^. Moreover, the CL precursor phosphatidylglycerol (PG) is enriched with the *n*-7 isomer in prostate cancer cell lines^25^. Our analysis confirmed these previous findings, and demonstrated that ELOVL5 knockdown or SCD1 inhibition reduced the relative abundance of the n-7 isomer in this phospholipid class. The detection of ^13^C_16_-palmitoleic acid in both PG and CL species in our prostate cancer cell lines indicates that ELOVL5 knockdown-induced changes in CL content may be controlled by *de novo* CL synthesis, as the labelled fatty acid is incorporated into both a precursor (PG) and the product (CL) (Fig. 4A). However, given the data presented in this study, we cannot discount the possibility that ELOVL5 targeting and reduced cVA production is merely altering diacyl and tetra-acyl speciation of PG and CL species in parallel. Interestingly, despite the small quantities of labelled palmitoleic and oleic acid supplemented relative to available fatty acids from culture media in these experiments, we were able to detect labelled cVA in CLs, indicating that it is being selectively and efficiently incorporated into this phospholipid species.

CLs are associated with many aspects of mitochondrial biology, including ROS regulation, cytochrome c-related cell death signalling and energy production^85–87^. Accordingly, we assessed mitochondrial ROS production and found that it was increased under ELOVL5 knockdown, and that exogenous cVA supplementation partially rescued this effect. When accounting for this effect, it is important to note that ELOVL5 knockdown not only reduces cVA abundance, it also increases the bioaccumulation of POA. POA is a well-defined lipokine^23^ and has been shown to effect aerobic glycolysis and lipid synthesis pathways^88^. Therefore, it is possible that the mitochondrial superoxide induction observed under ELOVL5 knockdown is being stimulated by the effects of POA, rather than the reduced cVA. At a minimum, we cannot distinguish between these two possibilities, however, the partial rescue of mitochondrial superoxide levels with cVA supplementation suggests that a combination of both effects is likely.

We have previously observed alterations to mitochondrial structure with ELOVL5 knockdown, including increased mitochondrial length and overall size^6^. Here, we demonstrate a change to mitochondrial cristae structure, specifically increased cristae number per mitochondrial surface, along with the changes to CL content under ELOVL5 knockdown. Altered cristae formation has previously been linked to cytochrome c release, where mutations in a protein that forms part of the mitochondrial contact site and cristae organizing system (MICOS) caused a more open cristae conformation that facilitated cytochrome c release^89^, however, we did not observe changes to cytochrome c abundance in the mitochondria. Cristae structures in the mitochondrial inner membrane are critical for ATP production via the ETC^86^, and CLs are required for efficient cristae packing and stabilisation of ETC complexes^58^. Furthermore, changes in CL content and cristae morphology within the IMM have previously been shown to influence mitochondrial ATP production^83,90,91^. Decreased CL content in skeletal muscle reduces ATP coupling efficiency (coupling of ATP synthase activity with ETC activity) in mice^83^, while similar reductions in overall CL content in glioblastoma stem cells were associated with decreased ETC and ATP production and reduced tumour growth^90^. In acute myeloid leukaemia cell lines, mitochondrial structural changes, including alterations to cristae conformation and density, controlled mitochondrial respiration^91^. To this effect, we have previously demonstrated that ELOVL5 protein level is positively correlated with mitochondrial respiratory capacity^6^. More specifically, the increased cristae density we observed with ELOVL5 knockdown could be linked to suboptimal ATP production, as the formation of long narrow cristae segments is thought to physically restrict diffusion of ADP into the cristae space, preventing its replenishment^92^. Furthermore, we found increased frequency and length of mitochondria-lipid droplet (LD) contacts in *ELOVL5* knockdown cells. The formation of these contacts is associated with mitochondrial stress^62,63^, whereby these tight organelle junctions can facilitate efficient transfer of fatty acids to maintain mitochondrial β-oxidation and ATP production during conditions of cellular starvation^61^. Taken together, these findings suggest that cVA incorporation in CLs can control mitochondrial energy homeostasis to regulate prostate cancer cell viability. Future investigations into specific alterations in mitochondrial ATP production, ETC complex activity and membrane stabilisation are warranted.

We suggest that the use of cVA over OA in the CLs of prostate cancer cell mitochondria represents a previously unrecognised method for fine tuning mitochondrial morphology and function. Through the incorporation of the less abundant cVA into CLs, prostate cancer cells can be selective about overall CL abundance and content, to optimise mitochondrial membrane properties associated with efficient ATP production. Moreover, cVA is likely to be predominantly sourced by *de novo* synthesis, as it is significantly less abundant than OA in the typical adult diet, found in dairy milk^93,94^ and sea buckthorn berries^95^, but is produced by ELOVL5 activity, which is enhanced in prostate cancer cells^6^, increasing the potential of the cell to control cVA availability and CL content. Our findings of similar uptake of cVA and OA by prostate cancer cells (Fig. S3D) support *de novo* cVA production by ELOVL5 as the primary mechanism to control cVA abundance, highlighting a potential therapeutic vulnerability of prostate cancer cells.

Studies on the effects of SCD1-produced MUFAs have been predominantly carried out in human cell lines or animal models^15,39–41^, while the reliance of clinical tumour cells on MUFAs was previously unaddressed. Our patient-derived explant (PDE) model recapitulates key features of native tumour cells, including the tumour microenvironment and response to oncogenic drivers^27^, providing a unique approach to study the effects of anti-cancer agents on individual patient samples. Of the nine PDEs tested here, only two did not respond to SCD1 inhibition, suggesting that MUFAs produced by SCD1 are important for optimal growth in prostate cancer tissue *ex vivo* and that several patients may benefit from targeted therapies. Broadly targeting MUFA production in cancer has already received significant attention and many SCD1 inhibitors have been designed and trialled in various cancer types^16,18,39,46,96–98^. Unfortunately, adverse side effects are often observed^18,98^, limiting their translation to the clinic. To reduce side effects, more specific targets are required. We suggest that pinpointing the mechanism of action of key MUFA species in specific cancer contexts could yield promising candidates. The data presented here suggests that prostate cancer cells have a unique reliance on cVA for cell growth and proliferation, over their benign counterparts. In addition, analysis of our clinical dataset found increased total lipid abundance and increased ratios of 18:1-to-16:1 abundance (representative of ELOVL5 activity) in certain lipid classes in patients that experienced clinical relapse compared to those that did not. Although preliminary, this finding is significant, as at the time the biopsies were taken, there was no histological differences between these tumours that pathologists could have used to distinguish the clinical outcomes of these patients. Given that clinical relapse and disease progression leads to the currently incurable and fatal form of prostate cancer, the potential to use ELOVL5 activity as a biomarker is an exciting prospect that warrants further investigation and validation.

## Acknowledgements

The authors thank the study participants, urologists, nurses, histopathologists, and Cassandra Gordon and Maurene Giles of the Australian Prostate Cancer Bioresource, who assisted in the recruitment and collection of patient material and information with assistance from Andrew Shepherd; Samira Khabbazi (The University of Adelaide, SA, Australia) for assistance with explant tissue collection and analysis; Swati Irani (The University of Adelaide, SA, Australia) and Madison Helm (The University of Adelaide, SA, Australia) for assistance with explant tissue culture and analysis; Nancy Santiapillai (The University of Sydney, NSW, Australia) for her technical input on the labelled fatty acid tracer lipidomics; and Sydney Mass Spectrometry, a core research facility at the University of Sydney, for providing the Mass Spectrometry instruments used in this study. JSS is supported by a University of Adelaide George Fraser Scholarship, a PhD Supplementary Scholarship from the Freemasons Centre for Male Health and Wellbeing, and a University of Adelaide SAiGENCI Short Term Scholarship. SJB acknowledges funding through the Australian Research Council (DP190101486) and generous support for mass spectrometry through the Central Analytical Research Facility (CARF) at the Queensland University of Technology. ZDN is supported by Cancer Australia (ID2011672) and the U.S. Department of Defense (PC210356). ZDN and LMB are supported by the Hospital Research Foundation (C-PJ-Prost-2020). LMB, AJH and JVS are supported by a Revolutionary Team Award from the Movember Foundation and the Prostate Cancer Foundation of Australia (MRTA3), and LMB, ML and JVS are supported by the U.S. Department of Defense (PC12714814).

## Author Contributions

JSS – investigation, formal analysis, writing – original draft, writing – review and editing; RSEY – investigation, formal analysis, writing – review and editing; LEQ - investigation, formal analysis, writing – review and editing; DCM – investigation, writing – review and editing; EE - investigation, formal analysis, writing – review and editing; JD – investigation, formal analysis; IRDJ – investigation; DAB – investigation; ML – conceptualisation, funding acquisition; AJH – writing – review and editing; SJB – writing – review and editing; JVS – conceptualisation, funding acquisition, writing – review and editing; ZDN – supervision, writing – review and editing; LMB – conceptualisation, funding acquisition, supervision, writing – review and editing.

