## Supplementary Figures for "*Cis*-vaccenic acid is a key product of stearoyl-CoA desaturase 1 and a critical oncogenic factor in prostate cancer"

### Supplementary Figure 1

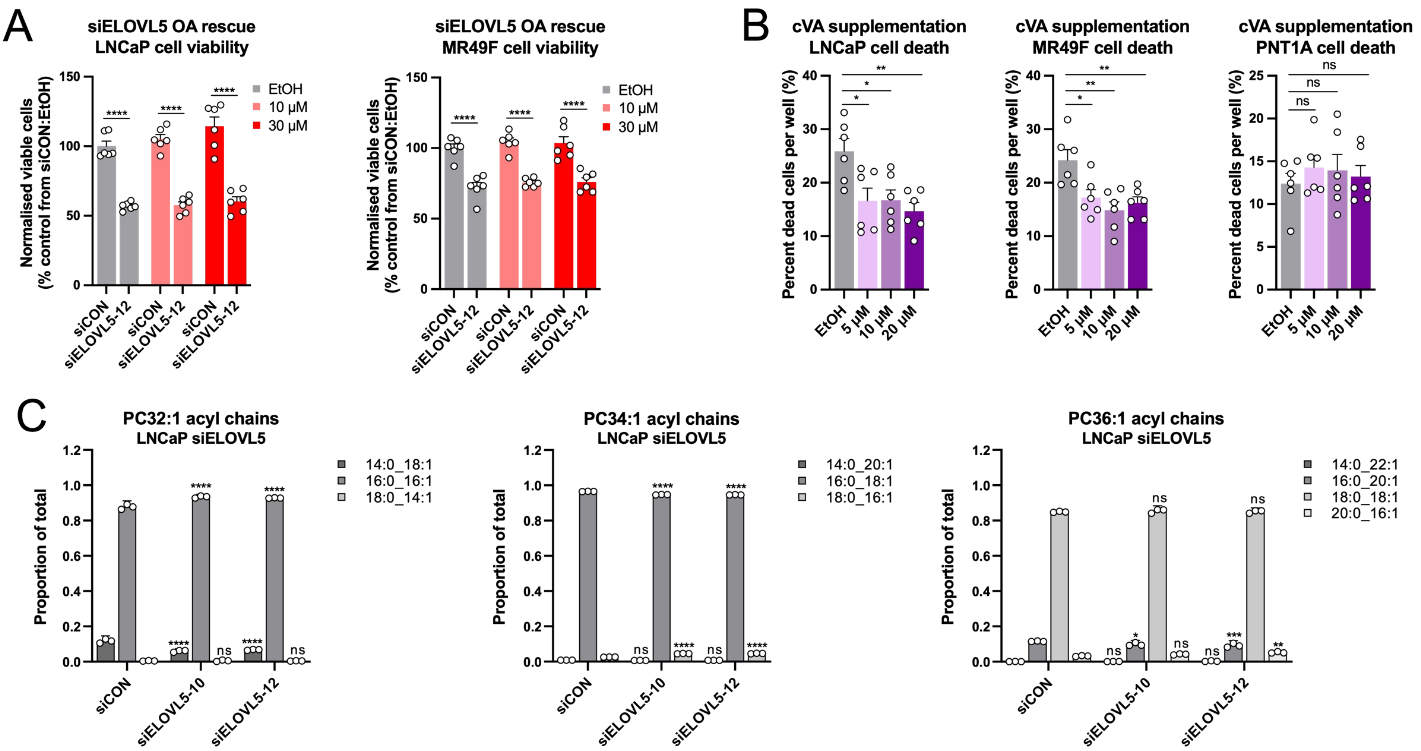

### Supplementary Figure 2

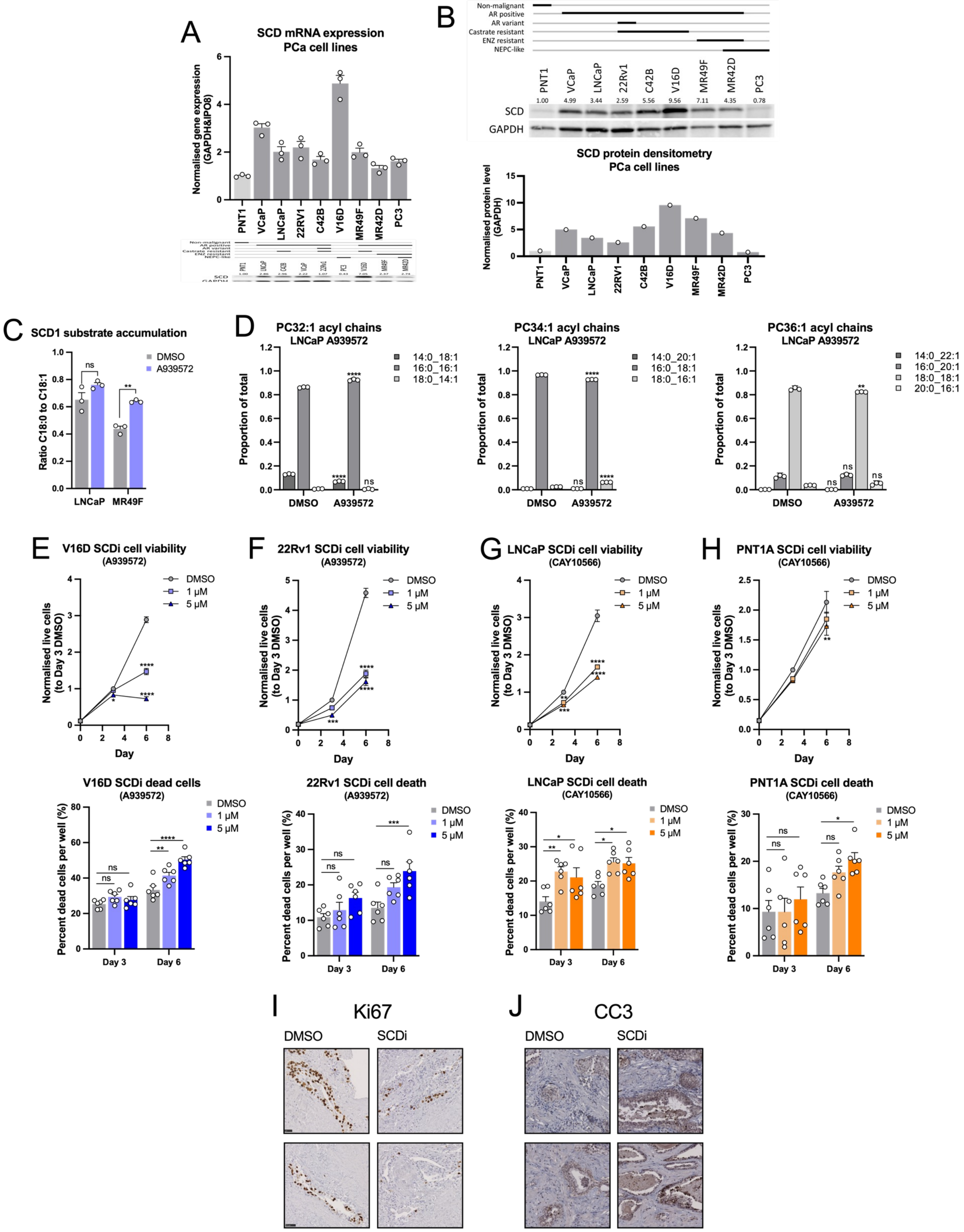

### Supplementary Figure 3

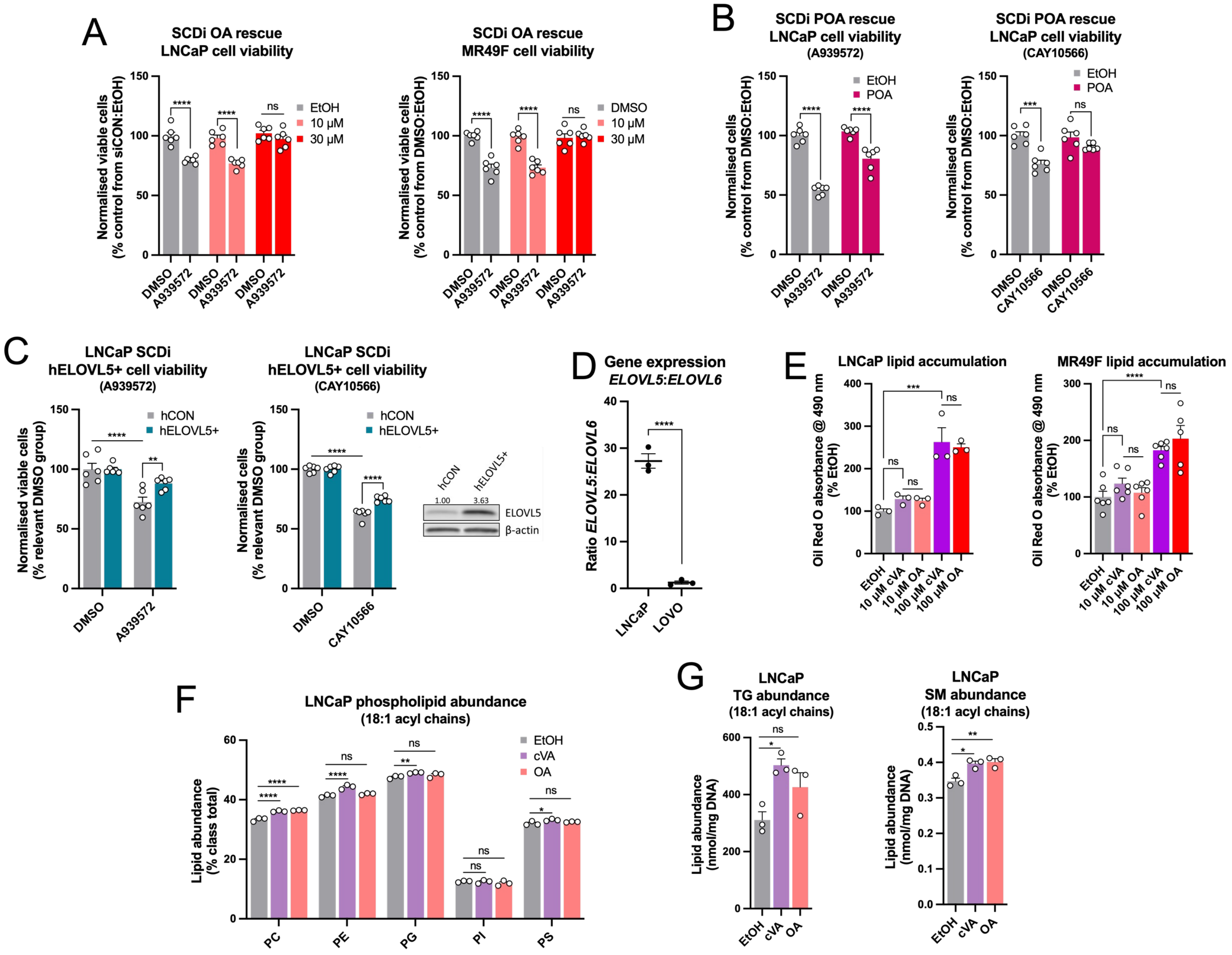

### Supplementary Figure 4

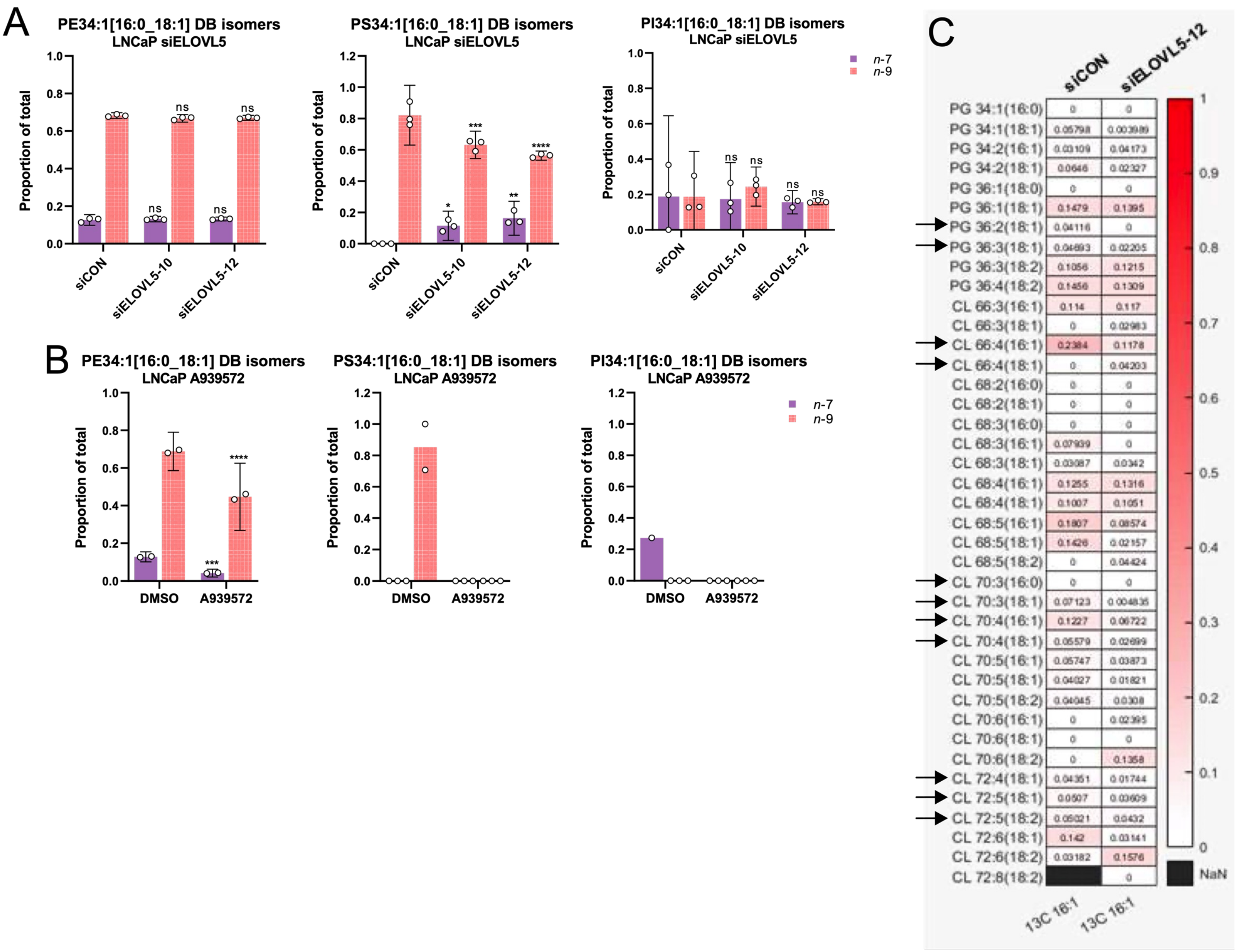

### Supplementary Figure 5

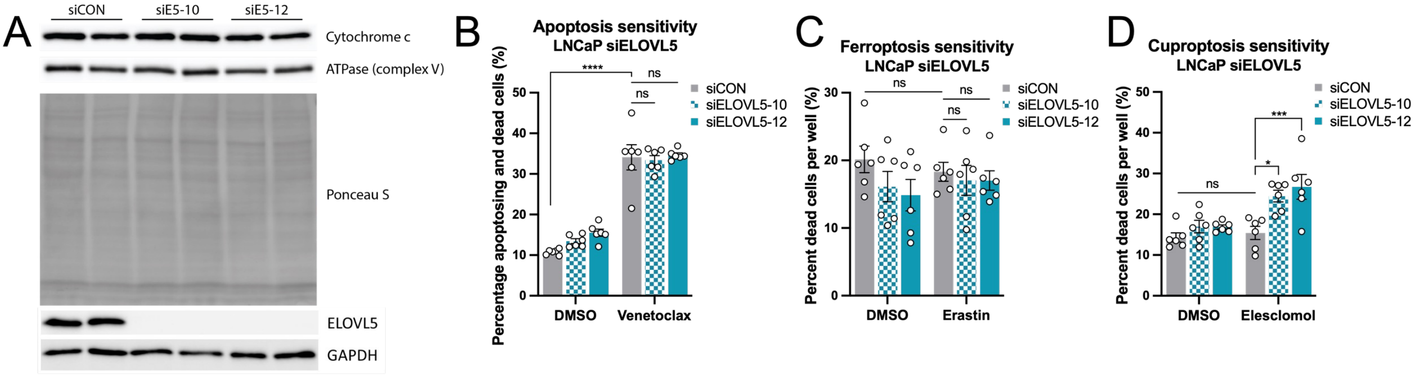

#### SUPPLEMENTARY FIGURE LEGENDS

##### Supplementary Figure 1.

[A] Cell viability experiments under ELOVL5 knockdown (siELOVL5) and supplementation of 10 or 30 μM oleic acid (OA) in prostate cancer cell lines LNCaP and MR49F. Data are presented as normalised mean ± SEM of two independent experiments (N = 2), with three technical replicates (n = 3). [B] Assessment of the effect of *cis*-vaccenic acid on cell death in prostate cancer cell lines LNCaP and MR49F, and the benign prostate cell line PNT1A. Data are presented as normalised mean ± SEM of two independent experiments (N = 2), with three technical replicates (n = 3). [C] Ozone-induced dissociation (OzID) lipidomic analysis of changes induced in acyl chain composition of phosphatidylcholines (PC) with compositions 32:1, 34:2 and 36:1 following ELOVL5 knockdown (siELOVL5). Data are presented as mean ± 95% confidence interval (CI) of one experiment (N = 1), with three technical replicates (n = 3). Statistical significance was determined using two-way ANOVA and multiple comparisons. * p<0.05; ** p<0.01; *** p<0.001; ****p<0.0001.

##### Supplementary Figure 2.

[A & B] Analysis of prostate cancer and benign cell lines for SCD1 mRNA expression [A] and protein levels [B]. Data are presented as mean ± SEM of one experiment. [C] Lipidomic analysis the ratio of C18:0 to C18:1 species in A939572-treated LNCaP cells. Data are presented as mean ± SEM of one experiment (N = 1), with three technical replicates (n = 3). [D] Ozone-induced dissociation (OzID) lipidomic analysis of changes induced in acyl chain composition of phosphatidylcholines (PC) with compositions 32:1, 34:2 and 36:1 in A939572-treated LNCaP cells. Data are presented as mean ± 95% confidence interval (CI) of one experiment (N = 1), with three technical replicates (n = 3). [E & F] Assessment of the effect of A939572 treatment on cell viability and cell death in prostate cancer cell lines V16D [E] and 22Rv1 [F]. Data are presented as normalised mean ± SEM of two independent experiments (N = 2), with three technical replicates (n = 3). [G & H] Assessment of the effect of CAY10566 treatment on prostate cancer cell line LNCaP [G] and the benign prostate cell line PNT1A [H]. Data are presented as normalised mean ± SEM of two independent experiments (N = 2), with three technical replicates (n = 3). [I] Representative images of Ki67 immunostaining of patient-derived explant samples treated with or without A939572 for 48 hours. [J] Representative images of cleaved caspase 3 immunostaining of patient-derived explant samples treated with or without A939572 for 48 hours. Statistical significance was determined using two-way ANOVA and multiple comparisons [C – H]. * p<0.05; ** p<0.01; *** p<0.001; ****p<0.0001.

##### Supplementary Figure 3.

[A] Cell viability rescue experiments in LNCaP and MR49F cells for A939572 treatment-related changes in response to 10 or 30 μM oleic acid (OA). Data are presented as normalised mean ± SEM of two independent experiments (N = 2), with three technical replicates (n = 3) in each. [B] Cell viability rescue experiments in LNCaP cells for A939572 or CAY10566 treatment-related changes in response to 10 μM palmitoleic acid (POA). Data are presented as normalised mean ± SEM of two independent experiments (N = 2), with three technical replicates (n = 3) in each. [C] LNCaP cells with stable ELOVL5 overexpression (hELOVL5+) were analysed for changes in cell viability with A939572 or CAY10566 treatment compared to control cells (hCON). Data are presented as normalised mean ± SEM of two independent experiments (N = 2), with three technical replicates (n = 3) in each. [D] Ratio of gene expression levels of *ELOVL5* and *ELOVL6* in LNCaP and LOVO cell lines. Data are presented as normalised mean ± SEM of one independent experiment (N = 1), with three technical replicates (n = 3). [E] Oil Red O analysis of lipid uptake in LNCaP and MR49F cells in response to 10 and 100 µM cVA or OA. Data are presented as normalised mean ± SEM of one independent experiment (N = 1), with three technical replicates (n = 3). [F] Lipidomic analysis of 18:1 acyl chains in phosphatidylcholine (PC), phosphatidylethanolamine (PE), phosphatidylglycerol (PG), phosphatidylinositol (PI) and phosphatidylserine (PS) in response to 10 µM cVA or OA. Data are presented as normalised mean ± SEM of one independent experiment (N = 1), with three technical replicates (n = 3). [G] Lipidomic analysis of 18:1 acyl chains in triacylglycerol (TG) and sphingomyelin (SM) lipid classes in response to 10 µM cVA or OA. Data are presented as mean ± SEM of one independent experiment (N = 1), with three technical replicates (n = 3). Statistical significance was determined by two-way ANOVA with multiple comparisons [A – C & F], student's unpaired t-test [D] or one-way ANOVA with multiple comparisons [E & G]. * p<0.05; ** p<0.01; *** p<0.001; ****p<0.0001.

**Supplementary Figure 4.**

[A & B] Ozone-induced dissociation (OzID) lipidomic analysis of changes induced in *n*-7 and *n*-9 isomer abundance in phosphatidylethanolamine (PE) 34:1, phosphatidylinositol (PI) 34:1 and phosphatidylserine (PS) 34:1 in ELOVL5 knockdown (siELOVL5) [A] and A939572-treated [B] LNCaP cells. Data are presented as mean ± 95% confidence interval (CI) of one experiment (N = 1), with three technical replicates (n = 3). [C] Lipidomic analysis of cardiolipin acyl chain composition in LNCaP ELOVL5 knockdown cells. Data are presented as mean ± SEM of one independent experiment (N = 1), with three technical replicates (n = 3). Statistical significance was determined by two-way ANOVA [A & B]. * p<0.05; ** p<0.01; *** p<0.001; ****p<0.0001.

**Supplementary Figure 5.**

[A] Western blot analysis of LNCaP ELOVL5 knockdown cells. Data are presented from one independent experiment (N = 1), with two technical replicates (n = 2). [B] Apoptosis measurement by flow cytometry for LNCaP ELOVL5 knockdown cells in response to 10 µM venetoclax treatment. Data are presented as mean ± SEM of two independent experiments (N = 2), with three technical replicates (n = 3) in each. [C] Cell death percentages of LNCaP ELOVL5 knockdown cells in response to 10 µM erastin treatment. Data are presented as normalised mean ± SEM of two independent experiments (N = 2), with three technical replicates (n = 3). [D] Cell death percentages of LNCaP ELOVL5 knockdown cells in response to 1 nM elesclomol treatment. Data are presented as normalised mean ± SEM of two independent experiments (N = 2), with three technical replicates (n = 3). Statistical significance was determined by two-way ANOVA [B – E]. * p<0.05; ** p<0.01; *** p<0.001; ****p<0.0001.
