## Supplementary Tables for "*Cis*-vaccenic acid is a key product of stearoyl-CoA desaturase 1 and a critical oncogenic factor in prostate cancer"

**Supplementary Table S1. Primary and secondary antibody details**

| Target | Company | Cat # | Species | Dilution | Block |
|---|---|---|---|---|---|
| Anti-Mouse biotinylated | Agilent, Dako | E0433 | Goat polyclonal | 1:1000 | 5% goat serum in 1 x PBS |
| Anti-Rabbit biotinylated | Agilent, Dako | E0432 | Goat polyclonal | 1:1000 | 5% goat serum in 1 x PBS |
| Anti-Mouse HRP | Agilent, Dako | P0161 | Rabbit polyclonal | 1:1000 | 3% skim milk in 0.1% TBS-Tween-20 |
| Anti-Rabbit HRP | Agilent, Dako | P0448 | Goat polyclonal | 1:1000 | 3% skim milk in 0.1% TBS-Tween-20 |
| Caspase 3 | Abcam | ab4051 | Rabbit polyclonal | 1:1000 | 5% goat serum in 1 x PBS |
| Cytochrome c | Abcam | ab90529 | Rabbit polyclonal | 1:1000 | 3% skim milk in 0.1% TBS-Tween-20 |
| ELOVL5 | Sigma-Aldrich | HPA047752 | Rabbit polyclonal | 1:2000 | 3% skim milk in 0.1% TBS-Tween-20 |
| GAPDH | Millipore | MAB374 | Mouse monoclonal | 1:5000 | 3% skim milk in 0.1% TBS-Tween-20 |
| Hsp90 | Cell Signalling Technology | 4874 | Rabbit polyclonal | 1:1000 | 3% skim milk in 0.1% TBS-Tween-20 |
| Ki67 | Agilent, Dako | M7240 | Mouse monoclonal | 1:500 | 5% goat serum in 1 x PBS |
| SCD | Sigma-Aldrich | HPA012107 | Rabbit polyclonal | 1:1000 | 3% skim milk in 0.1% TBS-Tween-20 |
| β-actin | Sigma-Aldrich | A5441 | Mouse monoclonal | 1:1000 | 3% skim milk in 0.1% TBS-Tween-20 |

**Supplementary Table S2. Primer sequences**

| Target gene | Forward sequence (5'-3') | Reverse sequence (5'-3') |
|---|---|---|
| ELOVL5 | GTGCACATTCCCTCTTGGTT | TTCAGGTGGTCTTTCCTTCG |
| GAPDH | TGCACCACCAACTGCTTAGC | GGCATGGACTGTGGTCATGAG |
| GUSB | CGTCCCACCTAGAATCTGCT | TTGCTCACAAAGGTCACAGG |
| IPO8 | GGCACCACTCAGCGAGGAT | AAGGTGAAGCCTCCCTGTTG |
| L19 | TGCCAGTGGAAAAATCAGCCA | CAAAGCAAATCTCGACACCTTG |
| SCD | ACACTTGGGAGCCCTGTATG | GACGATGAGCTCCTGCTGTT |

**Supplementary Table S3. Imaris Creation Parameters**

| **Function** | **Parameter Group** | **Parameter** |
|---|---|---|
| Surface (mitochondria) | [Algorithm] | Enable Region Of Interest = false |
| | | Enable Region Growing = false |
| | | Enable Tracking = false |
| | | Enable Shortest Distance = true |
| | [Source Channel] | Source Channel Index = 2 |
| | | Enable Smooth = true |
| | | Surface Grain Size = 0.200 µm |
| | | Enable Eliminate Backgroud = true |
| | | Diameter Of Largest Sphere = 0.250 µm |
| | [Threshold] | Enable Automatic Threshold = true |
| | | Manual Threshold Value = 113.138 |
| | | Active Threshold = true |
| | | Enable Active Threshold B = true |
| | | Manual Threshold Value B = 1016.48 |
| | | Active Threshold B = false |
| | [Classify Surface] | "Number of Voxels Img=1" above 10.0 |
| Spots (lipid droplets) | [Algorithm] | Enable Region Of Interest = false |
| | | Enable Region Growing = false |
| | | Enable Tracking = false |
| | | Enable Region Growing = false |
| | | Enable Shortest Distance = true |
| | [Source Channel] | Source Channel Index = 1 |
| | | Enable Smooth = true |
| | | Estimated XY Diameter = 0.700 µm |
| | | Estimated Z Diameter = 1.40 µm |
| | | Background Subtraction = true |
| | [Classify Surface] | "Intensity Max Ch=1 Img=1" above 1000 |
